# Immunity-longevity tradeoff neurally controlled by GABAergic transcription factor PITX1/UNC-30

**DOI:** 10.1101/2021.02.25.432801

**Authors:** Benson Otarigho, Alejandro Aballay

## Abstract

A body of evidence indicates that metazoan immune and aging pathways are largely interconnected, but the mechanisms involved in their homeostatic control remain unclear. In this study, we found that the PITX (paired like homeodomain) transcription factor UNC-30 controls the tradeoff between immunity and longevity from the nervous system in *Caenorhabditis elegans*. PITX/UNC-30 functional loss enhanced immunity in a GATA/ELT-2- and p38 MAPK/PMK-1-dependent manner and reduced longevity by activating MXD/MDL-1 and the C2H2-type zinc finger transcription factor PQM-1. The immune inhibitory and longevity stimulatory functions of PITX/UNC-30 required the sensory neuron ASG and a neurotransmitter signaling pathway controlled by NPR-1, which is a G protein-coupled receptor related to mammalian neuropeptide Y receptors. Our findings uncovered a suppressive role of GABAergic signaling in the neural control of a biological tradeoff where energy is allocated towards immunity at the expense of longevity.

## Introduction

The activation of an immunological response is an essential and costly energetic physiological process that results in the reallocation of resources, causing changes in the mechanisms that control general somatic maintenance (Ayres, 2020; Ganeshan and Chawla, 2014; Ganeshan et al., 2019; Jung et al., 2019). A tightly controlled immune activation is critical as its dysregulation has a major impact on a number of vital biological functions (Bird, 2019; Ganeshan et al., 2019; Jung et al., 2019; Kuchroo et al., 2012; Libert et al., 2006). In addition, the significant metabolic challenge posed by immune activation, immunopathology, and immune homeostasis necessitates an allocation of nutrients and energy that may lead to shortened longevity (Ayres, 2020; Backhed et al., 2019; Franceschi et al., 2018; Kaiser et al., 2020; Riera and Dillin, 2015; Solana et al., 2006; Tracey, 2002; Wani et al., 2019; Wu et al., 2019). Hence, the imbalance caused by immune activation results in a tradeoff between immunity and longevity that is evolutionarily conserved throughout the animal kingdom.

While the aforementioned changes to maintain homeostasis suggest the existence of a tradeoff between immunity and longevity across metazoans, the specific pathways involved in those tradeoffs and the underlying mechanisms that control them remain unknown. Several conserved pathways, including the DAF-2 (homolog of insulin-like growth factor 1, IGF1), AGE-1 (homolog of phosphatidylinositol 3-kinase, PI3K), DAF-16 (homolog of FOXO transcription factor), and HFS-1 (homolog of heat shock transcription factor 1, HSF1) pathways, control both immunity and longevity in the nematode *Caenorhabditis elegans* (Chávez et al., 2007; Garsin et al., 2003; Hajdu-Cronin et al., 2004; Libina et al., 2003; Mohri-Shiomi and Garsin, 2008; Morris et al., 1996; Murphy et al., 2003; Singh and Aballay, 2006). However, whether and how these interconnected immune and longevity pathways are regulated by tradeoff mechanisms still needs to be determined. The nervous system, which can integrate different cues in milliseconds and interpret conflicting stimuli, is well-equipped to effectively regulate tradeoffs.

To investigate the role of the nervous system in the control of the immunity-longevity tradeoff, we studied the GABAergic transcription factor PITX/UNC-30 in *C. elegans*. PITX/UNC-30 transcriptionally regulates the expression of GAD/UNC-25 and SLC32A1/UNC-47 that are critical in the biosynthesis, packaging, and trafficking of GABA (Cinar et al., 2005; Eastman et al., 1999b; Garcia et al., 2007; Gendrel et al., 2016; McIntire et al., 1997; Mclntire et al., 1993a; Mclntire et al., 1993b). We found that UNC-30 loss-of-function mutations led to enhanced resistance to killing by Gram-negative and Gram-positive bacterial pathogens. Conversely, UNC-30 loss-of-function mutations resulted in reduced longevity. The resistance to pathogen infection was mediated by the GATA transcription factor ELT-2 and the p38 mitogen-activated protein kinase PMK-1. The reduced longevity phenotype of UNC-30 mutants was due to higher activity of the MAX dimerization protein MXD/MDL-1 and the C2H2-type zinc finger transcription factor PQM-1. We further found that neuropeptide signaling mediated by the NPYR/NPR-1 pathway acts downstream of UNC-30 and the amphid sensory neuron ASG to control the tradeoff. Our findings highlight a neuronal network that controls the tradeoff between immunity and longevity. Given the conservation of GABAergic signaling and the longevity and immune pathways involved in the tradeoff, our findings raise the possibility that a similar process may occur in higher metazoans, including humans.

## Results

### Neuronal UNC-30 controls the immunity-longevity tradeoff

As a first step to understand the role of UNC-30 in the control of the immunity-longevity tradeoff, we studied the survival of *unc-30* loss-of-function animals infected with the human opportunistic pathogen *P. aeruginosa* strain PA14. We found that *unc-30(ok613)* mutants exhibited enhanced survival against *P. aeruginosa-*mediated killing compared to wild-type animals (Figure 1A). We also found that the *unc-30(ok613)* mutants exhibited less visible bacterial colonization and significantly reduced colony-forming units compared to wild-type animals (Figure 1B-C). Different *unc-30* strains also displayed enhanced survival against *P. aeruginosa* compared to wild-type animals (Figure S1A). When exposed to *E. coli*, which is the food source of *C. elegans* in the laboratory, *unc-30(ok613)* mutants exhibited enhanced survival compared to wild-type animals (Figure 1D and S1B). Because proliferating live *E. coli* is a cause of death in *C. elegans* and animals deficient in the immune response are persistently colonized and killed by the bacterium (Garigan et al., 2002; Otarigho and Aballay, 2020; Sutphin and Kaeberlein, 2009), we first investigated the survival of *unc-30(ok613)* animals on killed *E. coli*. Because animals fed heat-killed *E. coli* exhibit bloated intestinal lumens that result in up-regulation of immune pathways (Singh and Aballay, 2019b), we exposed the animals to lawns of *E. coli* killed by ultraviolet light (UV) that does not cause any intestinal distension (S2A-J). As shown in Figure 1E, the longevity of *unc-30(ok613)* animals is shorter than that of wild-type animals when exposed to UV-killed *E. coli*. As ampicillin was added to prevent the growth of any bacteria that might have escaped the heat treatment, we studied whether the addition of the antibiotic would impact the lifespan of the animals. As shown in Figure S1B, ampicillin did not affect the lifespan of *unc-30(ok613)* mutants.

**Figure 1.**
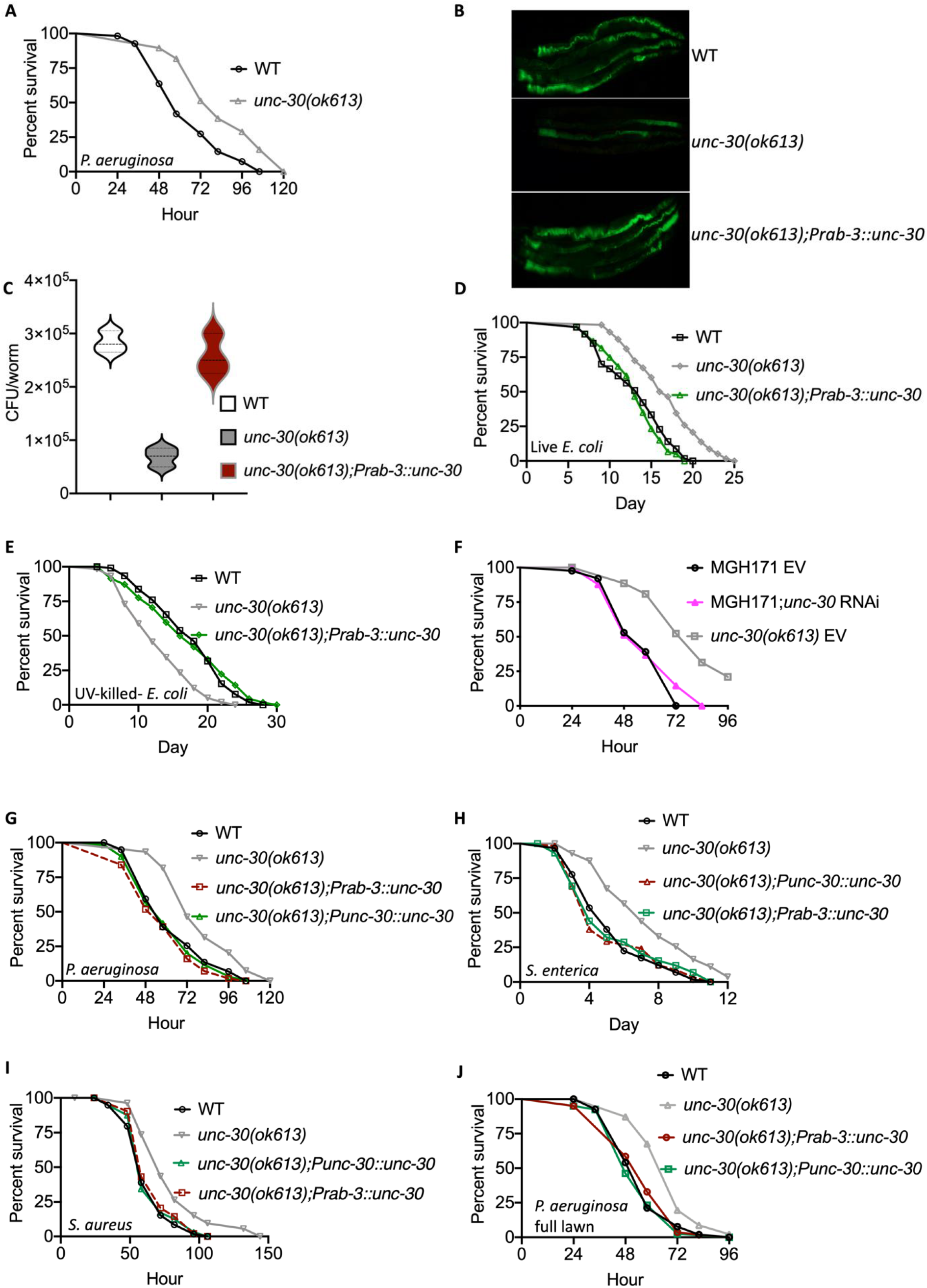
UNC-30 Functional Loss exhibits enhanced immunity and reduced longevity. (A) Wild type (WT) and *unc-30(ok613)* animals were exposed to *P. aeruginosa* partial lawn and scored for survival. WT vs *unc-30(ok613)*, P < 0.0001. (B) Colonization of WT, *unc-30(ok613)*, and *unc-30(ok613);Prab-3::unc-30* animals by *P. aeruginosa*-GFP after 24 hours at 25°C. (C) Colony-forming units per animal [WT, *unc-30(ok613)*, and *unc-30(ok613);Prab-3::unc-30*] grown on *P. aeruginosa* -GFP for 24 hours at 25°C. Bars represent means while error bars indicate SD; ****p < 0.001. (D) WT, *unc-30(ok613)*, and *unc-30(ok613);Prab-3::unc-30* animals were exposed live *E. coli* and scored for survival. WT vs *unc-30(ok613)*, P < 0.0001; *unc-30(ok613);Prab-3::unc-30*, P=NS. (E) WT, *unc-30(ok613)*, and *unc-30(ok613);Prab-3::unc-30* animals were exposed UV-killed *E. coli* and scored for survival. WT vs *unc-30(ok613)*, P < 0.0001; *unc-30(ok613);Prab-3::unc-30*, P=NS. (F) RNAi intestine-specific strain MGH171, MGH171; *unc-30* RNAi, and *unc-30(ok613)* animals were exposed to *P. aeruginosa* and scored for survival. EV, empty vector RNAi control. MGH171 EV vs. *unc-30(ok613)* EV, P < 0.0001; MGH171;*unc-30* RNAi, P = NS. (G) WT, *unc-30(ok613), unc-30(ok613);Punc-30::unc-30*, and *unc-30(ok613);Prab-3::unc-30* animals were exposed to *P. aeruginosa* partial lawn and scored for survival. WT vs *unc-30(ok613)*, P < 0.0001; *unc-30(ok613);Punc-30::unc-30*, and *unc-30(ok613);Prab-3::unc-30*, P = NS. (H) WT, *unc-30(ok613), unc-30(ok613);Punc-30::unc-30*, and *unc-30(ok613);Prab-3::unc-30* animals were exposed *S. enterica* partial lawn and scored for survival. WT vs *unc-30(ok613)*, P < 0.0001; *unc-30(ok613);Punc-30::unc-30*, and *unc-30(ok613);Prab-3::unc-30*, P = NS. (I) WT, *unc-30(ok613), unc-30(ok613);Punc-30::unc-30*, and *unc-30(ok613);Prab-3::unc-30* animals were exposed *S. aureus* partial lawn and scored for survival. WT vs *unc-30(ok613)*, P < 0.0001; *unc-30(ok613);Punc-30::unc-30*, and *unc-30(ok613);Prab-3::unc-30*, P = NS. (J) WT, *unc-30(ok613), unc-30(ok613);Punc-30::unc-30*, and *unc-30(ok613);Prab-3::unc-30* animals were exposed to *P. aeruginosa* full lawn and scored for survival. WT vs *unc-30(ok613)*, P < 0.0001; *unc-30(ok613);Punc-30::unc-30*, and *unc-30(ok613);Prab-3::unc-30*, P = NS.

UNC-30 is mainly expressed in three sensory neuronal cells (ASG, PVD, and OLL), three interneurons (PVP, RID, and RIH), and two motor neurons (DD and VD), as well as in the intestine (Cinar et al., 2005; Eastman et al., 1999b; Jin et al., 1994; Kurup and Jin, 2016; Walton et al., 2015; Westmoreland et al., 2001). To study whether UNC-30 may act cell autonomously in the intestine to control immune pathways, we knocked down UNC-30 in the intestine of an RNAi intestine-specific strain, MGH171. The lack of a significant difference between control and *unc-30* RNAi animals (Figure 1F) suggests that *unc-30* does not function in the intestine to control immunity and that it may control the immunity-longevity tradeoff from the nervous system. To study whether UNC-30 controls the immunity-longevity tradeoff in a cell non-autonomous manner from the nervous system, we rescued *unc-30* under the control of a pan-neuronal promoter and exposed the animals to *P. aeruginosa*. As a control, *unc-30* was also expressed using its native promoter. The results obtained show that UNC-30 expression driven by its own or the pan-neuronal promoter rescued both the enhanced survival of *unc-30(ok613)* animals to *P. aeruginosa*-mediated killing (Figure 1G) and the reduced longevity of the animals grown on heat-killed *E. coli* (Figure 1E). To study the specificity of pathogen susceptibility of *unc-30(ok613)* animals, we infected them with other human bacterial pathogens, including the Gram-negative bacterium *Salmonella enterica* strain 1344 and the Gram-positive bacteria *Staphylococcus aureus* strain NCTCB325. Loss of UNC-30 enhanced the nematode’s survival against all bacterial pathogens studied (Figure 1H-I), suggesting that UNC-30 suppresses *C. elegans* general defense against bacterial pathogens. Because *C. elegans* naturally exhibits an avoidance behavior when exposed to *P. aeruginosa*, which can be observed in conditions in which the animals can freely enter and exit the bacterial lawn, we used plates that were completely covered by bacteria to control for avoidance (full-lawn). The survival of *unc-30(ok613)* animals was also significantly higher than that of wild-type and *unc-30* rescued animals (Figure 1J). We also observed similar resistance to pathogen-mediated killing when the animals were exposed to full-lawns of *S. enterica* or *S. aureus* (Figure S3A-B). The neural expression of *unc-30* rescued the survival defect of *unc-30(ok613)* animals (Figures 1G-J). Consistent with the idea that the resistance of *unc-30(ok613)* animals is not due to enhanced pathogen avoidance, we found that the lawn occupancy of *unc-30(ok613)* mutants is comparable to that of wild-type animals (Figure S3C).

*C. elegans* feeds on bacteria and the pharyngeal contraction/pumping is a direct measure of food intake (Cao et al., 2017; Sellegounder et al., 2019; Singh and Aballay, 2019c; Styer et al., 2008). Thus, we asked whether the resistance to infections and reduction of the lifespan of *unc-30(ok613)* animals could be due to a reduction in pathogen intake. We, therefore, measured the pharyngeal pumping rates of *unc-30(ok613)* and wild-type animals on *P. aeruginosa* bacterial lawns as well as on heat-killed *E. coli*. We found that *unc-30(ok613)* animals exhibited pumping rates comparable to that of wild-type animals (Figure S3D), indicating that the dose of pathogens is similar in both cases. It has recently been demonstrated that bacterial accumulation in animals defective in the defecation motor program (DMP) causes intestinal distension that elicits a robust immune response (Singh and Aballay, 2019c) and modulates longevity (Kumar et al., 2019). We found that the defecation cycle of *unc-30(ok613)* animals is indistinguishable from that of wild-type animals (Figure S3E). Certain gene mutations in *C. elegans* affect the reproductive system, which could lead to reduced progeny or cause total sterility, resulting in increased resistance to pathogen infection (Berman and Kenyon, 2006; Powell and Ausubel, 2008). In addition, the tradeoff between immunity and longevity can be dependent on the reproductive dynamics of some mutant animals (Amrit et al., 2019). Thus, we compared the brood size of *unc-30(ok613)* and wild-type animals and did not observe any difference in the numbers of neither laid eggs nor brood size (Figure S4A-B). Taken together, these results suggest that *unc-30* functions in the nervous system to control a tradeoff between immunity and longevity.

### UNC-30 inhibits the expression of immune and age-related genes

To dissect the UNC-30-dependent immune and longevity mechanisms involved in the control of defense against pathogen exposure and longevity, we employed transcriptomic analyses to identify dysregulated immune genes and pathways in uninfected and *P. aeruginosa*-infected *unc-30(ok613)* and wild-type animals (Table S1). To identify gene groups that were controlled by UNC-30, we performed an unbiased gene enrichment analysis using a Wormbase enrichment analysis tool, https://wormbase.org/tools/enrichment/tea/tea.cgi, (Angeles-Albores et al., 2016), that is specific for *C. elegans* gene data analyses. The 10 gene ontology clusters with the highest enrichment score of vital biological functions for upregulated and downregulated genes in both non-infected and infected groups are shown in Figure 2A and S5A-B. Overall, the gene expression data showed that the most enriched and highly significant upregulated genes were a part of neuropeptide signaling and immune/defense response pathways in both non-infected and infected groups (Figure 2A and Table S2). Other genes linked to biological processes such as metabolism, response to biotic stimuli, and IRE1-mediated unfolded protein response, were also upregulated in *unc-30(ok613)* animals. Consistently with the role of UNC-30 in the control of longevity (Figure 1E), genes involved in aging were also significantly enriched (Figure 2A).

**Figure 2.**
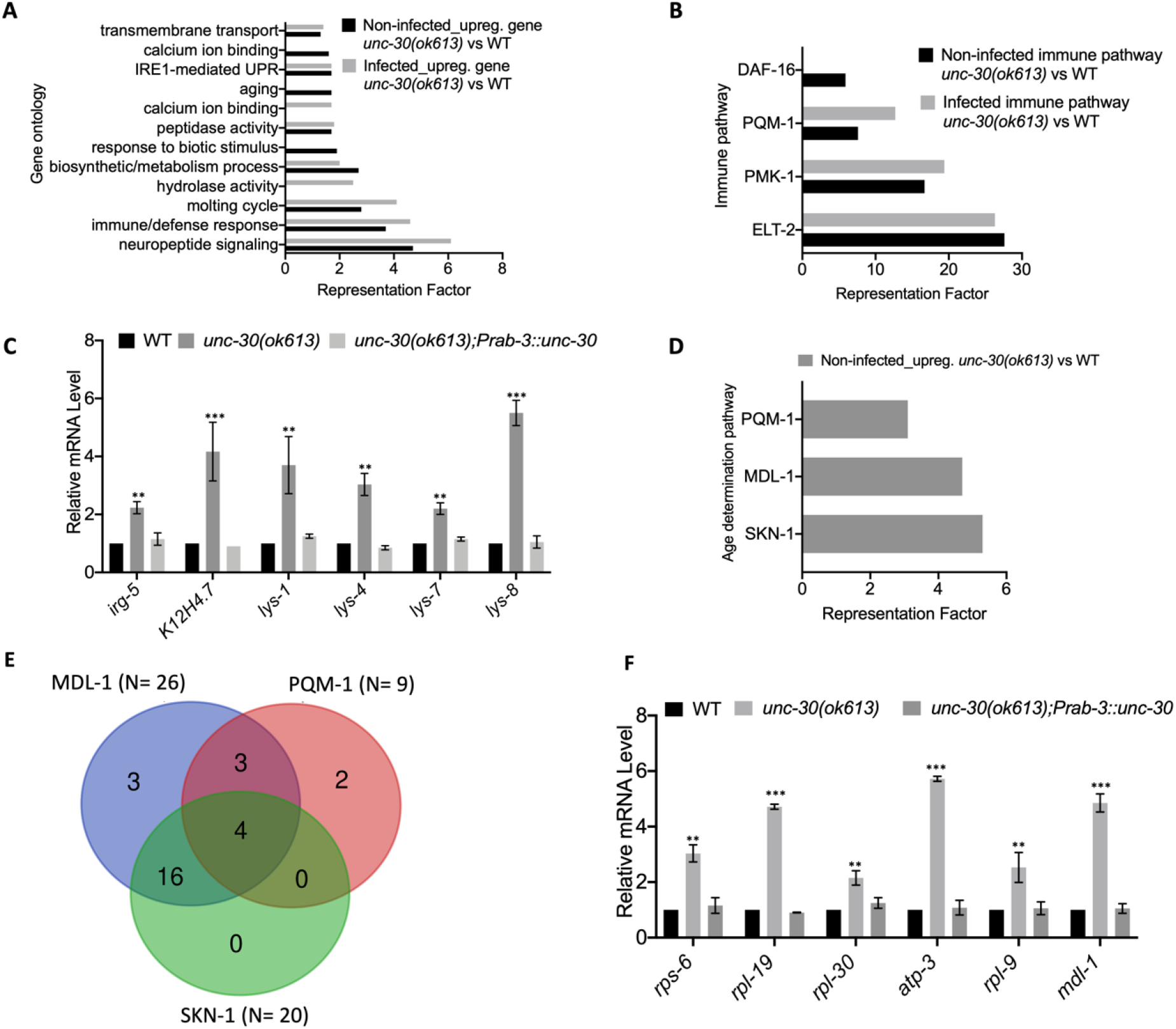
PITX1/UNC-30 regulates immune and age determination pathways. (A) Gene ontology analysis of upregulated genes in *unc-30(ok613)* vs WT in both non-infected and *P. aeruginosa*-infected animals. The cutoff is based on the filtering thresholds of P < 0.05 and arranged according to the representation factor. (B) Representation factors of immune pathways for the upregulated immune genes in *unc-30(ok613)* vs WT in both non-infected and *P. aeruginosa*-infected animals. (C) qRT-PCR analysis of immune gene expression in WT and *unc-30(ok613)* animals. Bars represent means while error bars indicate SD; *p < 0.05, **p < 0.001 and ***p < 0.0001. (D) Representation factors of age determination pathways for the upregulated aging genes in *unc-30(ok613)* vs WT in non-infected animals. (E) Venn diagram showing the age determination genes in each pathway for the upregulated aging genes in *unc-30(ok613)* vs WT in non-infected animals. (F) qRT-PCR analysis of age determination genes expression in WT and *unc-30(ok613)* animals. Bars represent means while error bars indicate SD; *p < 0.05, **p < 0.001 and ***p < 0.0001.

To identify the potential immune pathways that may play a role in the pathogen resistance to the UNC-30-deficient animals, we performed a gene enrichment analysis using the immune gene subset by employing the Worm Exp tool (https://wormexp.zoologie.uni-kiel.de/wormexp/) (Yang et al., 2016), which integrates all published expression datasets for *C. elegans* to analyze and generate pathways. We found that pathways required for *C. elegans* defense against bacterial infections were highly enriched, including the ELT-2, PMK-1, PQM-1, and DAF-2/DAF-16 insulin pathways (Figure 2B and Table S3) (Aballay et al., 2003; Garsin et al., 2003; Head et al., 2017; Huffman et al., 2004; Kerry et al., 2006; Kim et al., 2002; Murphy et al., 2003; Rajan et al., 2019; Shapira et al., 2006; Singh and Aballay, 2006, 2009; Troemel et al., 2006). We also noticed that several of the PMK-1 and DAF-16-dependent genes were also controlled by ELT-2 (Figure S6 and Tables S3), indicating that ELT-2-dependent genes may play a major role in the enhanced resistance to pathogen phenotype of *unc-30(ok613)* animals. Thus, we confirmed the role of neural UNC-30 in the control of ELT-2-dependent immune genes, including *lys-1, K12H4*.*7, lys-4, lys-7*, and *lys-8* (Figure 2C). These results indicate that UNC-30 regulates *C. elegans* defense against bacterial infections mainly by activating immune genes, several of which are controlled by the ELT-2 immune transcription factor.

We further employed the Worm Exp tool (https://wormexp.zoologie.uni-kiel.de/wormexp/) (Yang et al., 2016) to analyze the age determination genes to identify specific pathways that could be transcriptionally regulated by UNC-30. We found that the SKN-1, MDL-1, and PQM-1 age determination pathways were highly enriched (Figure 2D and Table S4) (Riesen et al., 2014; Tepper et al., 2013a; Tepper et al., 2014; Tullet et al., 2008). Although the SKN-1 cluster had the highest representation factor (Figure 2D), most of the genes up-regulated in *unc-30(ok613)* animals were under the control of the MDL-1 pathway (Figure 2E). Because the evolutionarily conserved transcription factors, MDL-1 and PQM-1 are known to suppress longevity and their inactivation results in lifespan extension (Riesen et al., 2014; Tepper et al., 2013b; Tepper et al., 2014), the upregulation of MDL-1 and PQM-1-dependent genes in *unc-30(ok613)* animals could explain the reduced longevity of the animals. We confirmed the role of UNC-30 in the control of MDL-1 and PQM-1-dependent genes (Figure 2F). In summary, the transcriptomic analyses revealed that while neural expression of UNC-30 downregulates the expression of immune genes controlled by ELT-2 and PMK-1, it upregulates age-related genes controlled by MDL-1 and PQM-1, indicating that UNC-30 controls a tradeoff between immunity and longevity.

### UNC-30 functional loss enhances immunity via ELT-2 and PMK-1 and reduces longevity via MDL-1 and PQM-1

To test the hypothesis that the enhanced resistance to *P. aeruginosa* infection of *unc-30(ok613)* animals is due to the upregulation of immune genes, we studied the role of suppression by RNAi of the immune pathways shown in Figure 2B. We inactivated *elt-2, pqm-1, pmk-1*, and *daf-16* in WT and *unc-30(ok613)* animals and exposed them to *P. aeruginosa*. Unlike *daf-16* and *pqm-1* RNAi (Figure 3A-B), *elt-2* RNAi was able to completely suppress the enhanced resistance to *P. aeruginosa* infection in *unc-30(ok613)* animals (Figure 3C). We also observed partial suppression of the enhanced pathogen resistance of *unc-30(ok613)* animals by *pmk-1* RNAi (Figure 3D). The lack of suppression of the phenotype of *unc-30(ok613)* animals by RNAi inhibition of *daf-16* and *pqm-1* was confirmed using *daf-16* and *pqm-1* mutant animals (Figure 3E-F). We also confirmed the partial suppression of the enhanced resistance to pathogen of *unc-30(ok613)* animals by inhibition of the PMK-1 pathway using *nsy-1(ag3)* and *sek-1(km4)* animals, which carry mutations in the genes encoding two kinases that function upstream PMK-1 (Figure 3G-H). These results indicate that loss of UNC-30 enhances immunity against infection by increasing the activity of ELT-2 and the PMK-1 pathway.

**Figure 3.**
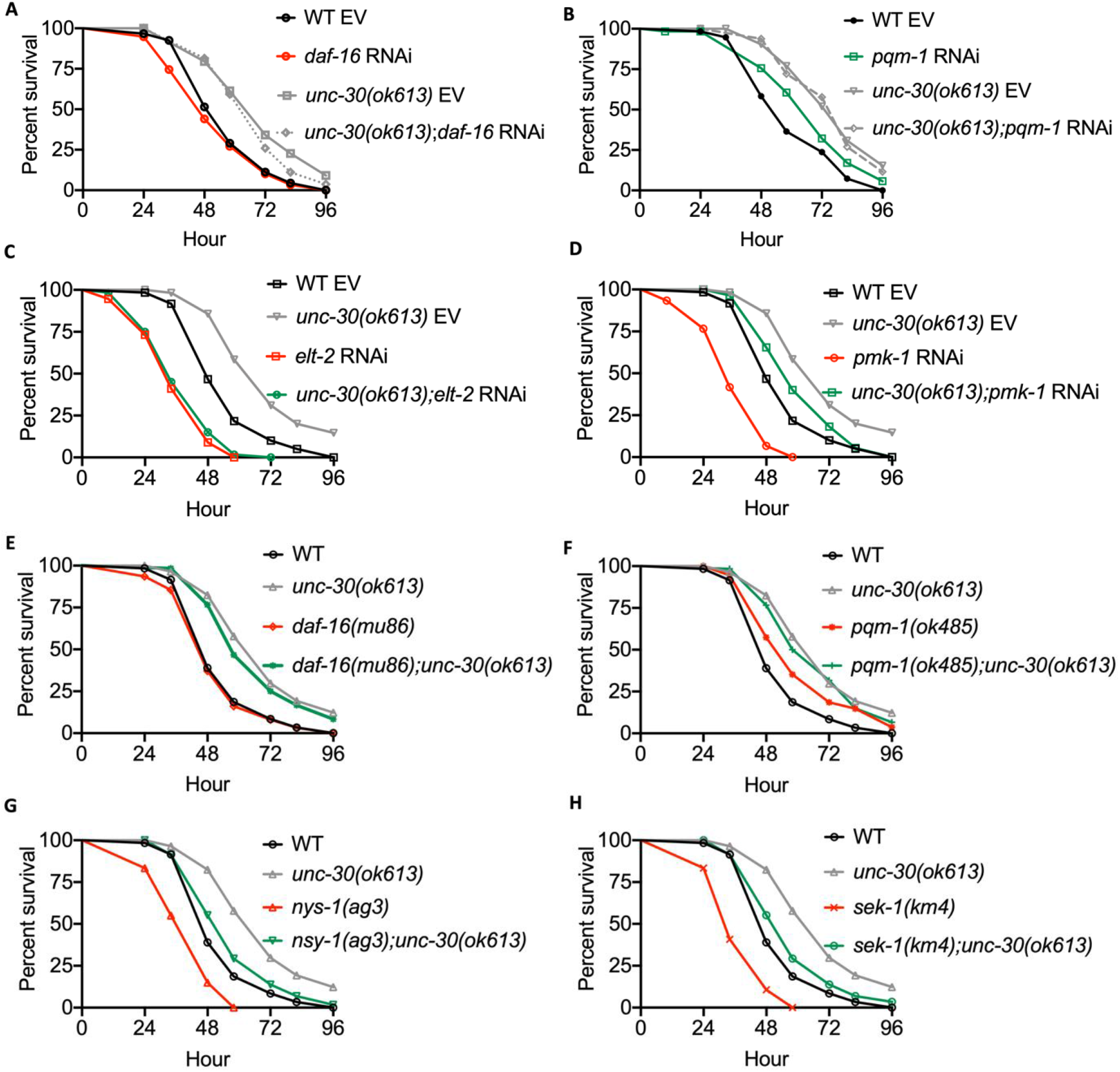
Functional loss of UNC-30 enhances immunity via the GATA/ELT-2 transcription factor and partly p38 MARK/PMK-1 pathway. (A) WT, *daf-16* RNAi, *unc-30(ok613)*, and *unc-30(ok613);daf-16* RNAi animals were exposed to *P. aeruginosa* and scored for survival. EV, empty vector RNAi control. WT EV vs. *unc-30(ok613)* EV, P < 0.0001; *daf-16* RNAi, P =NS; *unc-30(ok613);daf-16* RNAi, P< 0.0001. (B) WT, *pqm-1* RNAi, *unc-30(ok613)*, and *unc-30(ok613);pqm-1* animals were exposed to *P. aeruginosa* and scored for survival. EV, empty vector RNAi control. WT EV vs. *unc-30(ok613)* EV, P < 0.0001; *pqm-1* RNAi, P <0.05; *unc-30(ok613);pqm-1* RNAi, P< 0.0001. (C) WT, *elt-2* RNAi, *unc-30(ok613)*, and *unc-30(ok613);elt-2* RNAi animals were exposed to *P. aeruginosa* and scored for survival. EV, empty vector RNAi control. WT EV vs. *unc-30(ok613)* EV, P < 0.0001; *elt-2* RNAi, P <0.0001; *unc-30(ok613);elt-2* RNAi, P< 0.0001. *elt-2* RNAi vs *unc-30(ok613) elt-2* RNAi, P=NS. (D) WT, *pmk-1* RNAi, *unc-30(ok613)*, and *unc-30(ok613);pmk-1* RNAi animals were exposed to *P. aeruginosa* and scored for survival. EV, empty vector RNAi control. WT EV vs. *unc-30(ok613)* EV, P < 0.0001; *pmk-1* RNAi, P <0.0001; *unc-30(ok613);pmk-1* RNAi, P< 0.05. *pmk-1* RNAi vs *unc-30(ok613) pmk-1* RNAi, P<0.0001. (E) WT, *daf-16(mu86), unc-30(ok613)*, and *daf-16(mu86);unc-30(ok613)* animals were exposed to *P. aeruginosa* and scored for survival. WT vs. *unc-30(ok613)*, P < 0.0001; *daf-16(mu86)*, P =NS; *daf-16(mu86);unc-30(ok613)*, P< 0.0001. (F) WT, *pqm-1(ok485), unc-30(ok613)*, and *pqm-1(ok485);unc-30(ok613)* animals were exposed to *P. aeruginosa* and scored for survival. WT vs. *unc-30(ok613)*, P < 0.0001; *pqm-1(ok485)*, P <0.05; *pqm-1(ok485);unc-30(ok613)*, P< 0.0001. (G) WT, *nsy-1(ag3), unc-30(ok613)*, and *nsy-1(ag3);unc-30(ok613)* animals were exposed to *P. aeruginosa* and scored for survival. WT vs. *unc-30(ok613)*, P < 0.0001; *nsy-1(ag3)*, P< 0.0001; *nsy-1(ag3);unc-30(ok613)*, P=NS. (H) WT, *sek-1(km4), unc-30(ok613)*, and *sek-1(km4);unc-30(ok613)* animals were exposed to *P. aeruginosa* and scored for survival. WT vs. *unc-30(ok613)*, P < 0.0001; *sek-1(km4)*, P< 0.0001; *sek-1(km4);unc-30(ok613)*, P=NS.

The Venn diagram shown in Figure 4A indicates a poor overlap between aging and immune genes, suggesting that the misregulation of different pathways causes the immune and longevity phenotypes of animals deficient in UNC-30. Thus, we reasoned that the upregulation of the age-related genes in un-infected *unc-30(ok613)* mutants compared with wild-type animals (Figure 2A) could be responsible for the reduced longevity of *unc-30(ok613)* animals. Consistent with this idea, a number of the age-related genes that are upregulated in *unc-30(ok613)* animals have been reported to extend lifespan when mutated or inactivated by RNAi (Figure 4B and Table S4). Moreover, the majority of age-related genes upregulated in *unc-30(ok613)* mutants compared with wild-type animals are controlled by MDL-1 and PQM-1 (Figure 2E), which are known to suppress longevity. Thus, we studied whether inactivation of MDL-1 or PQM-1 could suppress the decreased longevity of *unc-30(ok613)* animals. As shown in Figures 4C and 4D, *mdl-1(tm311)* and *pqm-1(ok485)* mutation suppressed the reduced lifespan of *unc-30(ok613)* animals. Consistent with the idea that higher activity of both MDL-1 and PQM-1 is responsible for the reduced lifespan of *unc-30(ok613)* animals, the lifespan of the double mutants *mdl-1(tm311);unc-30(ok613)* and *pqm-1(ok485);unc-30(ok613)* is shorter than that of the single mutants *mdl-1(tm311)* and *pqm-1(ok485)*, respectively (Figures 4C and 4D). To study whether the longevity of the single mutants is greater than that of the double mutants due to higher PQM-1 or MDL-1 activity, we inhibited both PQM-1 and MDL-1 in *unc-30(ok613)* animals. As shown in Figure 4E, the longevity of *mdl-1* RNAi animals is not significantly different than that of *pqm-1(ok485);unc-30(ok613);mdl-1* RNAi animals. Inhibition of SKN-1 by RNAi reduced the longevity of wild-type animals but had no effect on *unc-30(ok613)* mutants (Figure 4F). This is expected because, unlike MDL-1 and PQM-1 that suppress longevity, SKN-1 promotes longevity. Taken together, these results indicate that neuronal UNC-30 suppresses ELT-2- and PMK-1 mediated immunity while prolonging longevity through the MDL-1 and PQM-1 pathways.

**Figure 4.**
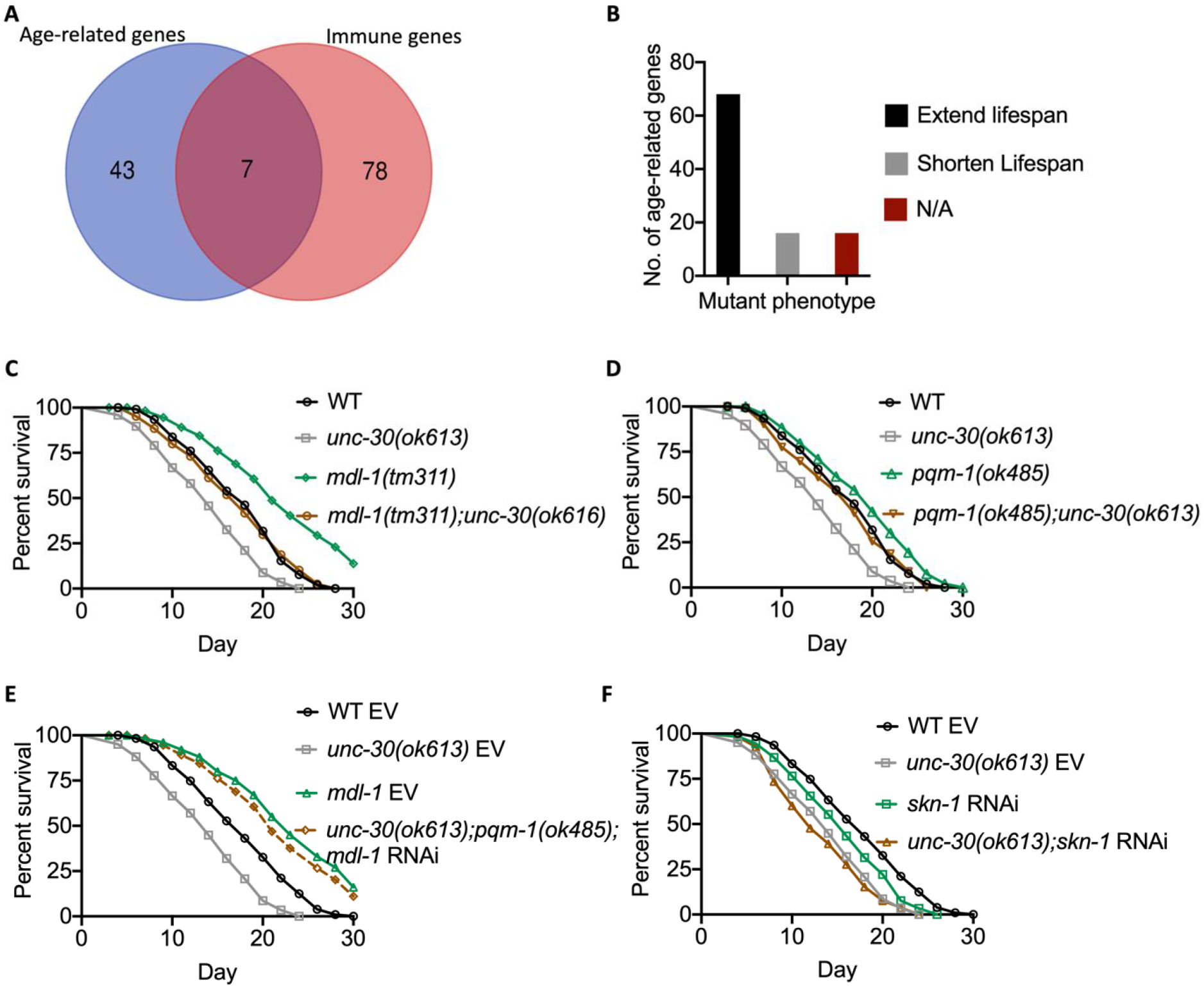
Functional loss of UNC-30 reduces longevity via MXD3/MDL-1 and PQM-1 pathways. (A) Venn diagram of the immune and aging genes upregulated in *unc-30(ok613)* vs WT non-infected animals. (B) Number of age determination genes upregulated in *unc-30(ok613)* vs WT for which mutant phenotypes have been reported. (C) WT, *mdl-1(tm311), unc-30(ok613)*, and *mdl-1(tm311);unc-30(ok613)* animals were exposed to UV-killed *E. coli* and scored for survival. WT vs. *unc-30(ok613)*, P < 0.0001; *mdl-1(tm311)*, P < 0.0001; *mdl-1(tm311);unc-30(ok613)*, P=NS. *mdl-1(tm311)* vs *mdl-1(tm311);unc-30(ok613)*, P < 0.001. (D) WT, *pqm-1(ok485), unc-30(ok613)*, and *pqm-1(ok485);unc-30(ok613)* animals were exposed to UV-killed *E. coli* and scored for survival. WT vs. *unc-30(ok613)*, P < 0.0001; *unc-30(ok613);pqm-1(ok485)*, P < 0.0001. *unc-30(ok613)* vs *unc-30(ok613);pqm-1(ok485)*, P< 0.05. (E) WT, *unc-30(ok613)*, and *unc-30(ok613);pqm-1(ok485);mdl-1* RNAi animals were exposed to UV-killed *E. coli* and scored for survival. EV, empty vector RNAi control. WT EV vs. *unc-30(ok613)* EV, P < 0.0001; *unc-30(ok613);pqm-1(ok485);mdl-1* RNAi, P<0.001. *unc-30(ok613);pqm-1(ok485);mdl-1* RNAi vs. *mdl-1* RNAi, P=NS. (F) WT, *skn-1* RNAi, *unc-30(ok613)*, and *unc-30(ok613);skn-1* RNAi animals were exposed to UV-killed *E. coli* and scored for survival. EV, empty vector RNAi control. WT EV vs *unc-30(ok613)* EV, P < 0.0001; *skn-1(mg570)/skn-1* RNAi, P < 0.001; *unc-30(ok613);skn-1* RNAi, P< 0.0001. *skn-1* RNAi vs *unc-30(ok613);skn-1* RNAi, P< 0.001.

### Neuronal UNC-30 regulates the immunity-longevity tradeoff via neuropeptide signaling

The subset of misregulated genes most significantly enriched in either infected or uninfected *unc-30(ok613)* animals corresponds to neuropeptide signaling genes (Figure 2A and Table S2). To identify potential neurotransmitter pathways that may be involved in the control of the tradeoff between immunity and longevity, we performed gene enrichment analysis using the neuropeptide genes subset (Table S5). Our findings show that most of the neuropeptide genes are transcriptionally controlled by NPR-1 (Figure 5A and Table S5). NPR-1 is a G protein-coupled receptor similar to neuropeptide Y receptor (NPYR) that controls *C. elegans* immune response against pathogens (Nakad et al., 2016; Reddy et al., 2009; Reddy et al., 2011; Singh and Aballay, 2019c; Styer et al., 2008). Other genes are controlled by miRNA and LIN-28 (Figure 5A and Table S5). Figure 5B shows confirmation of the misregulation of NPYR/NPR-1-dependent genes in *unc-30(ok613)* mutants compared to wild-type animals. We also found that the pan-neuronal expression of *unc-30(ok613)* rescued the changes in expression of NPR-1-dependent neuropeptide genes (Figure 5B.

**Figure 5.**
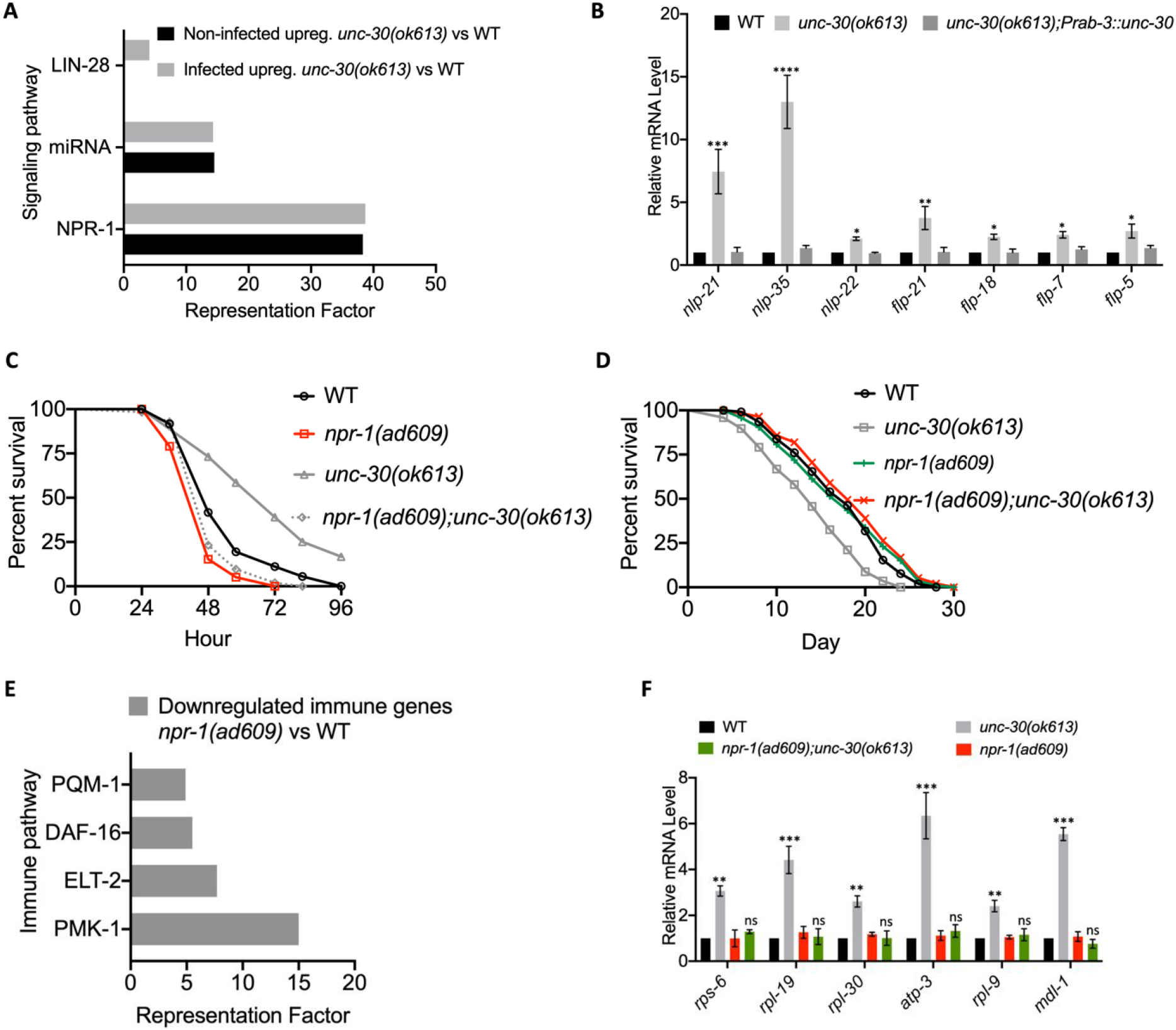
PITX1/UNC-30 regulates immunity and longevity via neuropeptide signaling. (A) Representation factors of neuropeptide genes upregulated in non-infected and *P. aeruginosa*-infected *unc-30(ok613)* vs WT animals. (B) qRT-PCR analysis of neuropeptide gene expression in WT, *unc-30(ok613)* and *unc-30(ok613);Prab-3::unc-30* animals. Bars represent means while error bars indicate SD; *p < 0.05, **p < 0.001 and ***p < 0.0001. (C) WT, *npr-1(ad609), unc-30(ok613)*, and *npr-1(ad609);unc-30(ok613)* animals were exposed to *P. aeruginosa* and scored for survival. WT vs *unc-30(ok613)*, P < 0.0001; *npr-1(ad609)*, P < 0.001; *npr-1(ad609);unc-30(ok613)*, P <0.05. *npr-1(ad609)* vs *npr-1(ad609)::unc-30(ok613)*, P = NS. (D) WT, *npr-1(ad609), unc-30(ok613)*, and *unc-30(ok613);npr-1(ad609)* animals were exposed to UV-killed *E. coli* and scored for survival. WT vs *unc-30(ok613)* P < 0.0001; *npr-1(ad609)*, P=NS. While *npr-1(ad609)* vs *unc-30(ok613)*, P< 0.0001; *unc-30(ok613);npr-1(ad609)*, P = NS. (E) Representation factors of immune genes downregulated in *npr-1(ad609)* vs WT. (F) qRT-PCR analysis of age determination gene expression in WT, *npr-1(ad609), unc-30(ok613)*, and *unc-30(ok613);npr-1(ad609)* animals animals. Bars represent means while error bars indicate SD; *p < 0.05, **p < 0.001 and ***p < 0.0001.

Because most *unc-30(ok613)* upregulated neuropeptide genes are NPYR/NPR-1-dependent, we asked whether NPR-1 may be part of the UNC-30 pathway that controls the immunity-longevity tradeoff. We hypothesized that if *npr-1* acts downstream of *unc-30, npr-1* mutation should suppress the enhanced immunity and reduced longevity of *unc-30(ok613)* animals. Indeed, we found that the *npr-1(ad609)* mutation was able to suppress the immunity and longevity phenotypes of *unc-30(ok613)* animals (Figure 5C, 5D, and S7). We further studied NPR-1-regulated genes and found that, in addition to controlling the immune PMK-1 pathway (Styer et al., 2008), NPR-1 controls immune genes that are ELT-2 and PQM-1-dependent (Figure 5E and Table S6). Consistent with the suppression of the reduced longevity of *unc-30(ok613)* animals by mutation in *npr-1*, we found that the *npr-1(ad609)* mutation suppressed the missregulation of age-related genes observed in *unc-30(ok613)* animals (Figure 5F).

### GABAergic ASG neurons control the UNC-30-modulated immunity-longevity tradeoff

Next, we asked which UNC-30-expressing neuronal cells could be involved in the control of the immune-longevity tradeoff. As a first step, we employed a two-component system to kill cells by using a reconstituted caspase (Chelur and Chalfie, 2007) to generate a PVP neuron ablated strain and that was exposed to *P. aeruginosa*. We focused on PVP because it is well connected to other UNC-30-expressing neurons, including DD and VD (Figure S8A). We found no significant difference between the PVP(-) and wild-type animals (Figure 6A), suggesting that the immunomodulatory function of UNC-30 is not linked to PVP neurons. However, we cannot rule out the role of DD, VD, and VC in immunity.

**Figure 6.**
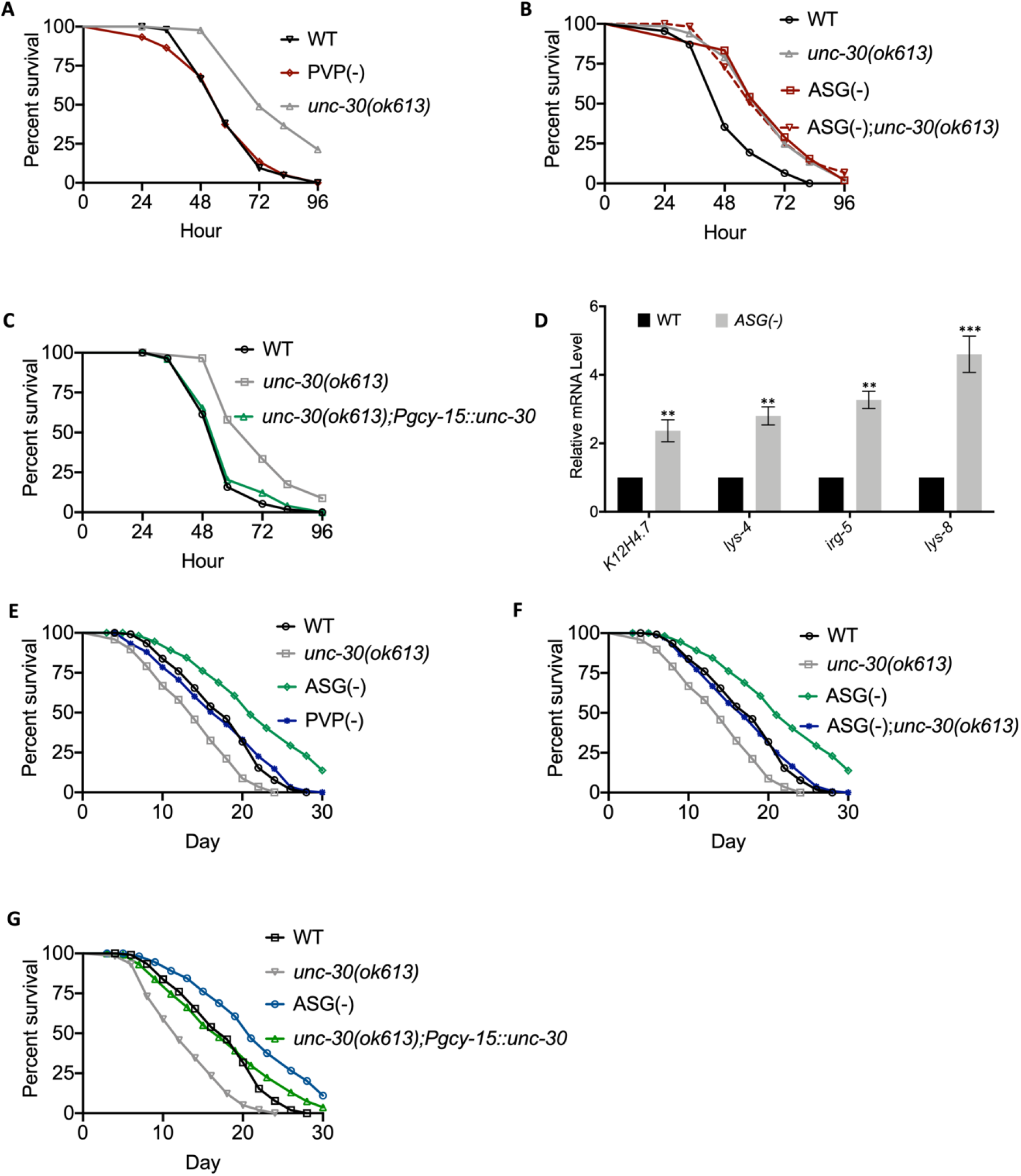
ASG neurons control the immunity-longevity tradeoff. (A) WT, PVP (-), and *unc-30(ok613)* animals were exposed to *P. aeruginosa* and scored for survival. WT vs PVP (-), P = NS. (B) WT, *unc-30(ok613)*, ASG(-), and ASG(-);*unc-30(ok613)* animals were exposed to *P. aeruginosa* and scored for survival. WT vs ASG(-), P < 0.0001; ASG(-)*;unc-30(ok613)*, P< 0.0001. While ASG(-) vs. ASG(-)*;unc-30(ok613)*, P = NS. (C) qRT-PCR analysis of immune gene expression in WT and ASG(-) animals. Bars represent means while error bars indicate SD; **p < 0.001 and ***p < 0.0001. (D) WT, *unc-30(ok613), unc-30(ok613);Pgyc-15::unc-30* animals were exposed to *P. aeruginosa* and scored for survival. WT vs *unc-30(ok613);Pgyc-15::unc-30*, P = NS. (E) WT, ASG(-), PVP(-), and *unc-30(ok613)* animals were exposed to UV-killed *E. coli* and scored for survival. WT vs ASG(-), P < 0.0001; *unc-30(ok613)*, P< 0.0001. (F) WT, ASG(-), ASG(-);*unc-30(ok613)*, and *unc-30(ok613)* animals were exposed to UV-killed *E. coli* and scored for survival. WT vs ASG(-), P < 0.0001; *unc-30(ok613)*, P< 0.0001. *unc-30(ok613)* vs ASG(-)::*unc-30(ok613)*, P < 0.001. (G) WT, *unc-30(ok613), unc-30(ok613);Pgyc-15::unc-30* animals were exposed to UV-killed *E. coli* and scored for survival. WT vs *unc-30(ok613);Pgyc-15::unc-30*, P = NS.

We next studied whether other UNC-30-expressing cells (ASG, PVD, or OLL sensory neurons) could control the defense response against pathogen infection. We employed an interactive tissue-gene expression prediction tool (http://worm.princeton.edu) to predict the expression enrichment level across different cells and tissues. The results obtained revealed that *unc-30* is highly expressed in ASG, which is an amphid neuron (Figure S8B) (Li and Kim, 2008; Pereira et al., 2015; Pocock and Hobert, 2010). Also, we noticed that some of the neuropeptide genes, including *flp-6, flp-13, flp-22*, and *ins-1*, that are upregulated in *unc-30(ok613)* animals are expressed in ASG neurons (Li and Kim, 2008; Pereira et al., 2015; Pocock and Hobert, 2010). Thus, we studied the effect of ASG ablation in the control of defense against *P. aeruginosa* infection in wild-type and *unc-30(ok613)* animals. We found that ASG(-) animals exhibited resistance to *P. aeruginosa*-mediated killing similar to that of *unc-30(ok613)* animals (Figure 6B). The resistance to *P. aeruginosa* infection of *unc-30(ok613)* and ASG(-)::*unc-30(ok613)* animals was also indistinguishable (Figure 6B). In addition, expression of *unc-30* only in ASG neurons completely suppressed the pathogen resistance of *unc-30(ok613)* animals (Figure 6C), suggesting that the GABAergic ASG neuron plays a key role in the UNC-30 modulation of immunity. To further confirm the role of ASG neurons in the control of immunity, we quantified the expression of immune genes and found that they were upregulated in ASG(-) animals (Figure 6D).

We also investigated the lifespan of animals lacking ASG(-) or PVP(-). As shown in Figure 6E, ASG(-) but not PVP(-) animals exhibited a longer lifespan than control animals. We also found that ASG ablation partially rescued the short lifespan of *unc-30(ok613)* animals, suggesting that ASG neurons participate in the control of longevity by UNC-30 (Figure 6F). We also found that *unc-30* expression in ASG neurons rescued the short lifespan of *unc-30(ok613)* animals (Figure 6G). In summary, our results suggest that neuronal UNC-30 and neuropeptide signaling regulate the tradeoff between immunity and longevity controlling the ELT-2 and PMK-1 immune pathway and the MDL-1 and PQM-1 aging pathways (Figure 7).

**Figure 7.**
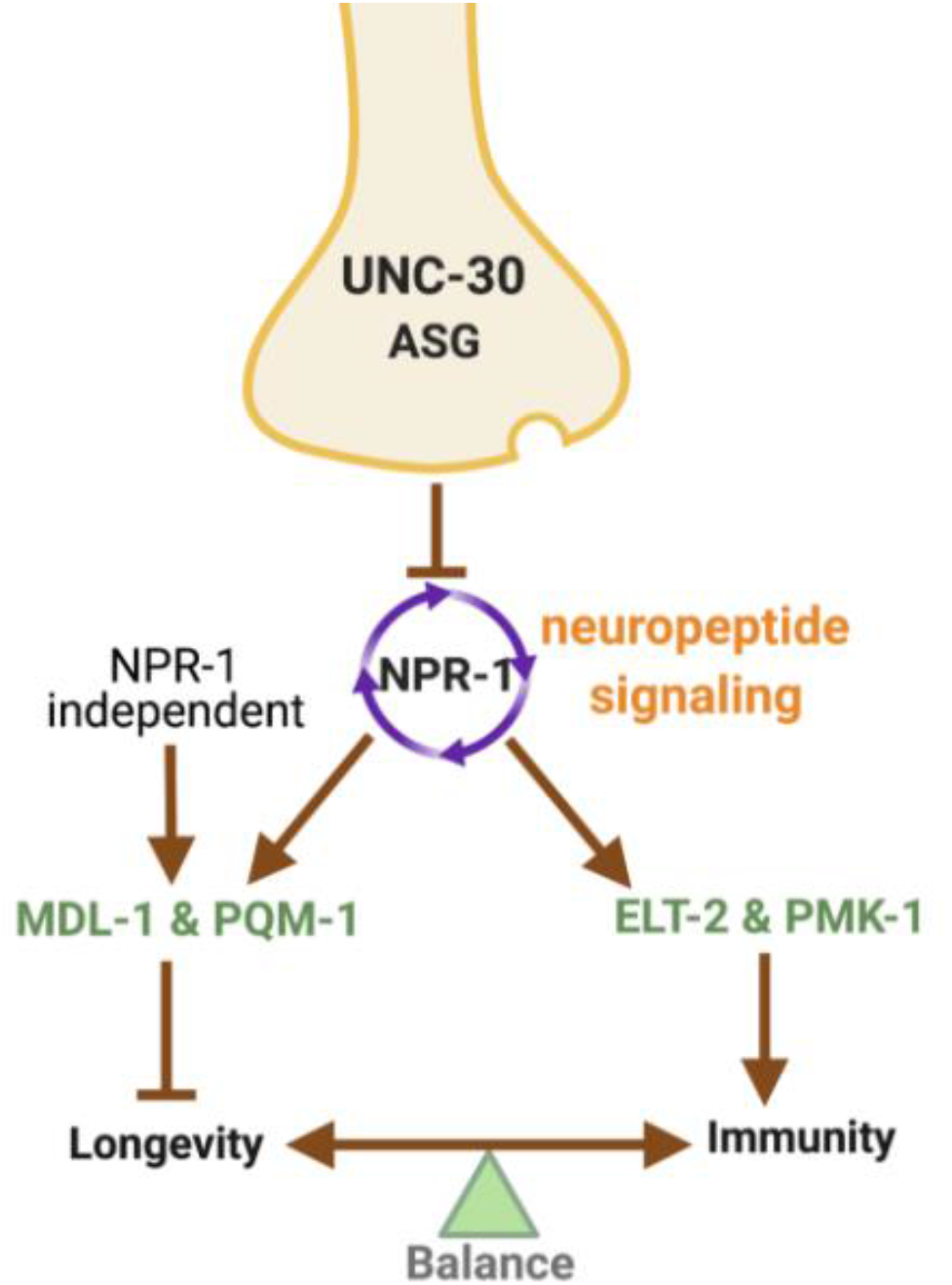
Model for neuronal control of the tradeoff between immunity and longevity via neuropeptide signaling in *C. elegans*. Neuronal PITX1/UNC-30, via the NPR-1, inhibits immunity by preventing the expression of ELT-2-and PMK-1-dependent immune genes and promotes longevity by preventing the expression of MDL-1 and PQM-1-dependent age-related genes.

## Discussion

Biological tradeoffs are widely employed by metazoans to maximize the allocation of precious resources that are critical for organismal homeostasis. However, the mechanisms that control them at the whole animal level remain obscure. The nervous system provides a perfect partner to help fine-tune tradeoffs because it can rapidly respond to many types of stimuli and is adept at controlling opposite functions. (Eastman et al., 1999a; Jin et al., 1994; Westmoreland et al., 2001). Here, we show that UNC-30 functions in the nervous system from where, together with neuropeptide signaling, it can control the tradeoff between immunity and longevity. The absence of UNC-30 results in the upregulation of the expression of age-related genes controlled by MDL-1 and PQM-1 and immune genes controlled by the ELT-2 and PMK-1. Furthermore, the GABAergic neuron ASG and the NPR-1 neuropeptide signaling are part of the UNC-30 pathway that controls the balance between immune activation and longevity.

Previous studies linked neurotransmitters such as dopamine and octopamine to the control of immunity via the PMK-1 pathway (Cao and Aballay, 2016; Pinoli et al., 2017; Sellegounder et al., 2019; Sun et al., 2011) and the X-box binding protein 1 (XBP-1) branch of the canonical UPR pathway (Cao et al., 2017). Acetylcholine has also been connected to the control of immunity via the intestinal epithelial Wnt Signaling (Labed et al., 2018). Furthermore, serotonin was also found to modulate the immune response via G-protein GOA-1 (Gαo) in rectal epithelial cells (Anderson et al., 2013; Hoffman and Aballay, 2019). Unlike other neurotransmitter pathways, the GABAergic signaling pathway modulates the immune response to pathogens via the control of the expression of immune genes regulated by ELT-2 and PMK-1. The UNC-30 pathway also controls genes that are involved in aging (Alcedo and Kenyon, 2004; Apfeld and Kenyon, 1999; Jeong et al., 2012; Kenyon, 2010; Wolkow et al., 2000). The G protein-coupled GABA receptor GBB-1 modulates longevity through G protein-PLCb, which transmits longevity signals to FOXO/DAF-16 (Chun et al., 2015; Yuan et al., 2019). Our studies highlight a role for neural UNC-30 in the integration of signals that control the balance between the activation of both immune and longevity pathways that have opposite effects. While UNC-30 suppresses immunity by inhibiting the expression of ELT-2- and PMK-1-dependent genes, it enhances longevity by inhibiting the activity of MDL-1 and PQM-1. Functional loss of MDL-1 and PQM-1 extends lifespan in *C. elegans* via DAF-2 insulin/IGF-1 (Johnson et al., 2014; Nakamura et al., 2016; Riesen et al., 2014; Templeman et al., 2018; Tepper et al., 2013b; Tepper et al., 2014). In addition, PQM-1 functions together with DAF-16 as a key transcriptional regulator of DAF-2-mediated longevity (Tepper et al., 2013b). However, the lack of significant enrichment in DAF-16-controled genes regulated by UNC-30 suggests that PQM-1 may also control longevity in a DAF-16-independent manner.

Our findings suggest that ASG neurons play a key role in controlling the tradeoff between immunity and longevity. They also express neuropeptides *flp-6, flp-13, flp-22*, and *ins-1* (Li and Kim, 2008; Pereira et al., 2015; Pocock and Hobert, 2010), which we found are upregulated in *unc-30(ok613)* animals. Furthermore, ASG cells respond to neuropeptides *flp-18* and *flp-21* via *npr-1* (Bargmann, 2006; Li and Kim, 2008; Pereira et al., 2015; Pocock and Hobert, 2010; Rex et al., 2005; Sugiura et al., 2005), which were upregulated in *unc-30(ok613)* animals. These results argue strongly that ASG neurons are critical in the control of the longevity-immunity tradeoff. Both PMK-1 and ELT-2 function in the intestine to control immunity (Bolz et al., 2010; Head et al., 2017; Kerry et al., 2006; Kim et al., 2002; Shapira et al., 2006). In addition, the role of PQM-1 and MDL-1 in longevity has been linked to the intestine (Riesen et al., 2014; Tepper et al., 2013b), suggesting that UNC-30 controls the immunity-longevity tradeoff in a cell non-autonomous manner. A recent example of a tradeoff mechanism indicates that dietary restriction extends lifespan by downregulating the PMK-1-mediated immune signaling (Wu et al., 2019). Further work will be required to establish whether this and other tradeoffs are controlled cell non-autonomously by the nervous system.

Our results provide evidence that specific genes and neurons in the nervous system act in a cell non-autonomous manner to coordinately control immune and aging pathways. Organisms widely rely on tradeoff mechanisms when a given trait cannot increase without decreasing another to maintain homeostasis. Given the evolutionary conservation of UNC-30 and the immune and aging pathways that are neutrally controlled, the identification and characterization of the cues that the nervous system uses to control the immunity-longevity tradeoff in *C. elegans* should yield insights into similar mechanisms used across metazoans.

## Materials and Methods

### Bacterial strains

The bacterial strains used in this study are *Escherichia coli* OP50, *E. coli* HT115(DE3) (Brenner, 1974), *Pseudomonas aeruginosa* PA14, *P. aeruginosa* PA14-GFP (Tan et al., 1999), *Salmonella enterica* Serovar Typhimurium 1344 (Wray and Sojka, 1978), and *Staphylococcus aureus* strain NCTCB325 (Sifri et al., 2003). Gram-negative bacteria were grown in Luria-Bertani (LB) broth. *Staphylococcus aureus* strain NCTCB325 was grown in Tryptic Soy Agar prepared with Nalidixic acid. All bacteria were grown at 37°C.

### *C. elegans* Strains and Growth Conditions

Hermaphrodite *C. elegans* (var. Bristol) wild type (N2) was used as control unless otherwise indicated. *C. elegans* strains CF1038 *daf-16(mu86)*, KU25 *pmk-1(km25)*, MGH171 *alxIs9* [*vha-6p::sid-1::SL2::GFP*], VC295 *unc-30(ok613)*, RB711 *pqm-1(ok485)*, DA609 *npr-1(ad609)*, CB318 *unc-30(e318)*, CB845 *unc-30(e191)*, SD39 *unc-30(e596)*, GR2245 *skn-1(mg570)*, RB711 *pqm-1(ok485)*, KU4 *sek-1(km4*) and AU3 *nsy-1(ag3)* were obtained from the Caenorhabditis Genetics Center (University of Minnesota, Minneapolis, MN). *mdl-1(tm311)* was obtained from National Bioresource Project (NBRP), Japan. *npr-1(ad609);unc-30(ok613), pqm-1(ok485);unc-30(ok613), daf-16(mu86);unc-30(ok613), sek-1(km4*)*;unc-30(ok613), nsy-1;unc-30(ok613), mdl-1(tm311);unc-30(ok613)*, and ASG(-)::*unc-30(ok613)* were obtained by standard genetic crosses. Rescued strain *unc-30(ok613);Punc-30::unc-30*, neuronal rescued strain *unc-30(ok613);Prab-3::unc-30* and *unc-30* rescued in ASG neuron-specific, *unc-30(ok613);Pgcy-15::unc-30* were generated as described below. The ASG(-) ablated Ex[Ptax-2::CZ::ced-3(p17)::unc-54 3’UTR + Plim-6::ced-3(p15)-NZ::unc-54 3’UTR, pRF4] and PVP(-) ablated Ex[Podr-2b::CZ::ced-3(p17)::unc-54 3’UTR, Punc-53::ced-3(p15)-NZ::unc-54 3’UTR, pRF4] were also generated as described below. The strains were crossed with the wild-type laboratory N2.

All strains were grown at 20^°^C on nematode growth medium (NGM) plates seeded with *E. coli* OP50 as the food source (Brenner, 1974) unless otherwise indicated. The recipe for the control NGM plates is: 3 g/l NaCl, 3 g/l peptone, 20 g/l agar, 5 μg/ml cholesterol, 1 mM MgSO4, 1 mM CaCl2, and 25 mM potassium phosphate buffer (pH 6.0). The NGM plates were without antibiotics except indicated.

### RNA Interference (RNAi)

Knockdown of targeted genes was obtained using RNAi by feeding the animal with *E. coli* strain HT115(DE3) expressing double-stranded RNA (dsRNA) homologous to a target gene (Fraser et al., 2000; Timmons and Fire, 1998). RNAi was carried out as described previously (Sun et al., 2011; Singh and Aballay, 2017). Briefly, *E. coli* with the appropriate vectors were grown in LB broth containing ampicillin (100 μg/mL) and tetracycline (12.5 μg/mL) at 37^°^C overnight and plated onto NGM plates containing 100 μg/mL ampicillin and 3 mM isopropyl β-D-thiogalactoside (IPTG) (RNAi plates). RNAi-expressing bacteria were grown at 37^°^C for 12-14 hours. Gravid adults were transferred to RNAi-expressing bacterial lawns and allowed to lay eggs for 2-3 hours. The gravid adults were removed, and the eggs were allowed to develop at 20^°^C to young adults. This was repeated for another generation (except for ELT-2 RNAi) before the animals were used in the experiments. The RNAi clones were from the Ahringer RNAi library.

### *C. elegans* Survival Assay on Bacterial Pathogens

*P. aeruginosa* and *S. enterica* were incubated in LB medium. *S. aureus* was incubated in TSA medium with nalidixic acid (10 μg/mL). The incubations were done at 37°C with gentle shaking for 12 hours. *P. aeruginosa* and *S. enterica* were grown on modified NGM agar medium (0.35% peptone). For partial lawn assays, 20 μl of the overnight bacterial cultures were seeded at the center of the relevant agar plates without spreading. For full lawn experiments, 20 μl of the bacterial culture were seeded and spread all over the surface of the agar plate. No antibiotic was used for *P. aeruginosa* and *S. enterica*, while nalidixic acid (10 μg/mL) was used for the TSA plates for *S. aureus*. The seeded plates were allowed to grow for 12 at 37°C. The plates were left at room temperature for at least 1 hr before the infection experiments. 20 synchronized young adult worms were transferred to the plates for infection, and three technical replicate plates were set up for each condition (n = 60 animals) and the experiments were performed in triplicate (Table 7). The plates were then incubated at 25°C. Scoring was performed every 12 hr for *P. aeruginosa* and *S. aureus*, while 24 hr for *S. enterica*. Worms were scored as dead if the animals did not respond to touch by a worm pick or lack of pharyngeal pumping. Live animals were transferred to fresh pathogen lawns each day. All *C. elegans* killing assays were performed three times independently.

### Bacterial lawn avoidance assay

Bacterial lawn avoidance assays were performed by 20 mL of *P. aeruginosa* PA14 on 3.5-cm modified NGM agar (0.35% peptone) plates and cultured at 37°C overnight to have a partial lawn. The modified NGM plates were left to cool to room temperature for about 1 hour, twenty young adult animals grown on *E. coli* OP50 were transferred to the center of each bacterial lawn after it. The number of animals on the bacterial lawns was counted at 12 and 24 hours after exposure.

### Pharyngeal pumping rate assay

Wild-type and *unc-30(ok613)* animals were synchronized by placing 20 gravid adult worms on NGM plates seeded with *E. coli* OP50 and allowing them to lay eggs for 60 min at 20°C. The gravid adult worms were then removed, and the eggs were allowed to hatch and grow at 20°C until they reached the young adult stage. The synchronized worms were transferred to NGM plates fully seeded with P. aeruginosa for 24 hours at 25°C. Worms were observed under the microscope with a focus on the pharynx. The number of contractions of the pharyngeal bulb was counted over 60 s. Counting was conducted in triplicate and averaged to obtain pumping rates.

### Defecation rate assay

Wild-type and mutant animals were synchronized by placing 20 gravid adult worms on NGM plates seeded with *E. coli* OP50 and allowing them to lay eggs for 60 min at 20°C. The gravid adult worms were then removed, and the eggs were allowed to hatch and grow at 20°C until they reached the young adult stage. The synchronized worms were transferred to NGM plates fully seeded with *P. aeruginosa* for 24 hours at 25°C. Worms were observed under a microscope at room temperature. For each worm, an average of 10 intervals between two defecation cycles was measured. The defecation cycle was identified as a peristaltic contraction beginning at the posterior body of the animal and propagating to the anterior part of the animal followed by feces expulsion.

### Brood Size Assay

The brood size assay was done following the earlier described methods (Kenyon et al., 1993; Otarigho and Aballay, 2020). Ten L4 animals from egg-synchronized populations were transferred to individual NGM plates (seeded with *E. coli* OP50) (described above) and incubated at 20°C. The animals were transferred to fresh plates every 24 hours. The progenies were counted and removed every day.

### *C. elegans* Longevity Assays

Longevity assays were performed on NGM plates containing live, heat killed, or UV-killed *E. coli* strains HT115 or OP50 as described earlier (Otarigho and Aballay, 2020; Sun et al., 2011)(Kumar et al., 2019; Sutphin and Kaeberlein, 2009). Animals were scored as alive, dead, or gone each day. Animals that failed to display touch-provoked or pharyngeal movement were scored as dead. Experimental groups contained 60 to 100 animals and the experiments were performed in triplicate (Table 8). The assays were performed at 20°C.

### Intestinal Bacterial Loads Visualization and Quantification

Intestinal bacterial loads were visualized and quantified as described earlier (Otarigho and Aballay, 2020; Sun et al., 2011). Briefly, *P. aeruginosa*-GFP lawns were prepared as described above. The plates were cooled to ambient temperature for at least an hour before seeding with young gravid adult hermaphroditic animals and the setup was placed at 25^°^C for 24 hours. The animals were transferred from *P. aeruginosa*-GFP plates to the center of fresh *E. coli* plates for 10 min to eliminate *P. aeruginosa*-GFP on their body. The step was repeated two times more to further eliminate external *P. aeruginosa*-GFP left from earlier steps. Subsequently, ten animals were collected and used for fluorescence imaging to visualize the bacterial load while another ten were transferred into 100 µL of PBS plus 0.01% Triton X-100 and ground. Serial dilutions of the lysates (10^1^-10^10^) were seeded onto LB plates containing 50 µg/mL of kanamycin to select for *P. aeruginosa*-GFP cells and grown overnight at 37 °C. Single colonies were counted the next day and represented as the number of bacterial cells or CFU per animal.

### Fluorescence and bloating Imaging

Fluorescence and bloating imaging was carried out as described previously (Otarigho and Aballay, 2020; Singh and Aballay, 2019a, c). Briefly, animals were anesthetized using an M9 salt solution containing 50 mM sodium azide and mounted onto 2% agar pads. The animals were then visualized for bacterial load using a Leica M165 FC fluorescence stereomicroscope. The diameter of the intestinal lumen was measured using Fiji-ImageJ software. At least 10 animals were used for each condition.

### RNA Sequencing and Bioinformatic Analyses

Approximately 40 gravid WT and *unc-30(ok613)* animals were placed for 3 hours on 10-cm NGM plates (seeded with *E. coli* OP50) (described above) to have a synchronized population, which developed and grew to L4 larval stage at 20^°^C. Animals were washed off the plates with M9 and frozen in QIAzol by ethanol/dry ice and stored at -80 prior to RNA extraction. Total RNA was extracted using the RNeasy Plus Universal Kit (Qiagen, Netherlands). Residual genomic DNA was removed using TURBO DNase (Life Technologies, Carlsbad, CA). A total of 6 μg of total RNA was reverse-transcribed with random primers using the High-Capacity cDNA Reverse Transcription Kit (Applied Biosystems, Foster City, CA).

The library construction and RNA sequencing in Illumina NovaSeq 6000 platform was done following the method described by (Zhu et al., 2018) and (Yao et al., 2018) pair-end reads of 150 bp were obtained for subsequent data analysis. The RNA sequence data were analyzed using a workflow constructed for Galaxy (https://usegalaxy.org) as described (Afgan et al., 2018; Afgan et al., 2016; Amrit and Ghazi, 2017). The RNA reads were aligned to the *C. elegans* genome (WS271) using the aligner STAR. Counts were normalized for sequencing depth and RNA composition across all samples. Differential gene expression analysis was then performed on normalized samples. Genes exhibiting at least two-fold change were considered differentially expressed. The differentially expressed genes were subjected SimpleMine tools from wormbase (https://www.wormbase.org/tools/mine/simplemine.cgi) to generate information such as wormBase ID and gene name, which are employed for further analyses. Gene ontology analysis was performed using the WormBase IDs in DAVID Bioinformatics Database (https://david.ncifcrf.gov) (Dennis et al., 2003) and validated using a *C. elegans* data enrichment analysis tool (https://wormbase.org/tools/enrichment/tea/tea.cgi). Immune pathways were obtained using the Worm Exp version 1 (http://wormexp.zoologie.uni-kiel.de/wormexp/)(Yang et al., 2016). The Venn diagrams were obtained using the web tool InteractiVenn (http://www.interactivenn.net) (Heberle et al., 2015) and bioinformatics and evolutionary genomics tool (http://bioinformatics.psb.ugent.be/webtools/Venn/). Tissue-gene expression prediction was used to simulate the expression pattern of UNC-30 (http://worm.princeton.edu) (Kaletsky et al., 2018). While neuron wiring was done using the database of synaptic connectivity of *C. elegans* for computation (White et al., 1986) http://ims.dse.ibaraki.ac.jp/ccep-tool/.

### RNA Isolation and Quantitative Reverse Transcription-PCR (qRT-PCR)

Animals were synchronized and total RNA extraction was done following the protocol described above. qRT-PCR was conducted using the Applied Biosystems One-Step Real-time PCR protocol using SYBR Green fluorescence (Applied Biosystems) on an Applied Biosystems 7900HT real-time PCR machine in 96-well-plate format. Twenty-five-microliter reactions were analyzed as outlined by the manufacturer (Applied Biosystems). The relative fold-changes of the transcripts were calculated using the comparative *CT*(2^-ΔΔ*CT*^) method and normalized to pan-actin (*act-1, -3, -4*). The cycle thresholds of the amplification were determined using StepOnePlus^™^ Real-Time PCR System Software v2.3 (Applied Biosystems). All samples were run in triplicate. The primer sequences were available upon request and presented in Table S9.

### Generation of transgenic *C. elegans*

To generate *unc-30* rescue animals, the *unc-30* DNA was amplified from the genomic DNA of Bristol N2 *C. elegans* adult worms. Plasmid pPD160_*Punc-30_unc-30* was constructed by linearization of plasmid pPD160_*Punc-30* using XbaI and SmaI restriction digestion enzymes. The amplified *unc-30* DNA was cloned behind its native promoter in the plasmid pPD160_*Punc-30*, between XbaI and SmaI sites. For the neuronal rescue strain, the plasmid pPD95_77_*Prab-3_unc-30* was constructed by cloning the amplified *unc-30* DNA into KpnI/HindIII digested pPD95_77_*Prab-3* under the promoter of *rab-3*. The constructs were purified and sequenced. Young adult hermaphrodite *unc-30(ok613) C. elegans* were transformed by microinjection of plasmids into the gonad as described (Mello and Fire, 1995; Mello et al., 1991). Briefly, a mixture containing pPD160_*Punc-30_unc-30* (40 ng/μl) and *Pmyo-3*::mCherry pCFJ104 (5 ng/μl) that drives the expression of mCherry to the muscle as a transformation marker were injected into the animals. For the neuronal rescue, a mixture containing pPD95_77_*Prab-3_unc-30* plasmids (40 ng/μl) and *Pmyo-2*::mCherry (5 ng/μl) pCFJ90 that drives the expression of mCherry to the pharynx as a transformation marker were injected into the animals.

The neuron ablated strains were generated by employing a two-component system reconstituted caspase (recCaspase) for selective ablation of targeted cells (Chelur and Chalfie, 2007). The ASG ablated neuron was done by separately cloning the promoters of *tax-2* and *lim-6* in the recCaspases plasmids. Briefly, a 1817bp segment of the *tax-2* promoter was cloned in the CZ-*ced-3*(p17):)::*unc-54* 3’UTR plasmid at the Sph I and EcoR I site to generate the *Ptax-2*::CZ::*ced-3*(p17)::*unc-54* 3’UTR plasmid, and a 5001bp segment of the *Lim-6* promoter was cloned in the *ced-3*(p15)+NZ:: *unc-54* 3’UTR plasmid at the Sph I and EcoR I site to generate the *Plim-6::ced-3*(p15)-NZ::*unc-54* 3’UTR plasmid. A mixture containing *Ptax-2-CZ-ced-3(p17)-unc-*54 3’UTR, *Plim-6-ced-3(p15)-NZ-unc*-54 3’UTR of 25ng/µL, and the roller marker pRF4 *rol-6(su1006)* (50ng/µL) was injected into the N2 young animals. To rescue *unc-30* in ASG neurons, *gcy-15* promoter (1987bp) and *unc-30* (5042bp) were amplified from gDNA. The *unc-30* fragment was cloned downstream of the *Pgcy-15* by in-fusion PCR to obtain *Pgcy-15::unc-30*, which was cloned into pPD95.77 between the Sph I and EcoR I sites to generate pPD95_77_*Pgcy-15::unc-30*. The ASG neuron–specific rescue strain *unc-30(ok613);Pgcy-15::unc-30* was generated by injecting plasmid pPD95_77_*Pgcy-15::unc-30* (25 ng/μl) with *Pmyo-2*::mCherry (10 ng/μl) as a co-injection marker into the *unc-30(ok613)* animals.

The PVP ablation was done by separately cloning the promoters of *odr-2* b isoform and *unc-53* in the recCaspases plasmids. Briefly, a 2661bp segment of the *odr-2* b isoform was cloned in the *CZ::ced-3(p17)::unc-*54 3’UTR plasmid at the Sph I and EcoR I site to obtain the *Podr-2b::CZ::ced-3(p17)::unc-54* 3’UTR plasmid and a segment of 3012bp of unc-53 was cloned in the *ced-3(p15)-NZ::unc-54* 3’UTR plasmid at the Sph I and EcoR I site to obtain the *Punc-53-ced-3(p15)-NZ-unc-54* 3’UTR plasmid. A mixture containing 25ng/µL of the *Podr-2b-CZ-ced-3(p17)-unc-54* 3’UTR plasmid, 25ng/µL of the *Punc-53-ced-3(p15)-NZ-unc-54* 3’UTR plasmid, and 50ng/µL of the roller marker pRF4 *rol-6(su1006)* was injected into N2 young animals. The successful transformation was determined by the identification of the selection marker. At least three independent lines carrying extrachromosomal arrays were obtained for each construct. Only worms with dominant roller phenotypes were selected for further experiment. Primers used in the generation of transgenic animals are shown in Table S9.

### Quantification and Statistical Analysis

Statistical analysis was performed with Prism 8 version 8.1.2 (GraphPad). All error bars represent the standard deviation (SD). The two-sample t test was used when needed, and the data were judged to be statistically significant when p < 0.05. In the figures, asterisks (*) denote statistical significance as follows: ns, not significant, *, p < 0.05, **, p < 0.001, ***, p < 0.0001, as compared with the appropriate controls. The Kaplan-Meier method was used to calculate the survival fractions, and statistical significance between survival curves was determined using the log-rank test. All experiments were performed at least three times.

## ACKNOWLEDGMENTS

This work was fully supported by NIH grants GM0709077 and AI117911 (to A.A.). Most strains used in this study were obtained from the Caenorhabditis Genetics Center (CGC), which is funded by the NIH Office of Research Infrastructure Programs (P40 OD010440) and the National BioResource Project (NBRP) of Japan.

## AUTHOR CONTRIBUTIONS

B.O and A.A. conceived and designed the experiments. B.O. performed the experiments. B.O. and A.A. analyzed the data and wrote the paper.

