## Supplemental Figures 1-8 for "Immunity-longevity tradeoff neurally controlled by GABAergic transcription factor PITX1/UNC-30"

**Supplemental Information**

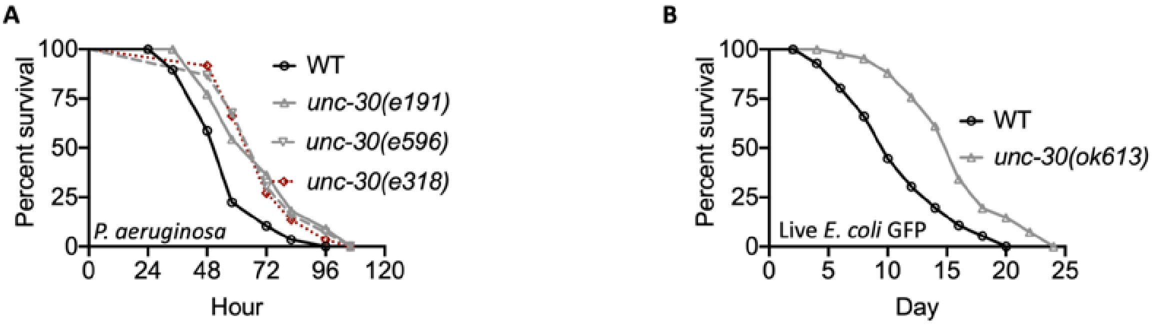


**Figure S1. UNC-30 functional loss enhances *C. elegans* pathogen-resistance and inhibits longevity. Related to Figure 1.**

(A) WT and *unc-30(e191), unc-30(e596),* and *unc-30(e318)* animals were exposed to *P. aeruginosa* partial lawn and scored for survival. WT vs *unc-30(e191); unc-30(e596)* and *unc-30(e318)*, $P < 0.0001$.

(B) WT and *unc-30(ok613)* animals were exposed live *E. coli* (OP50 GFP resistance to ampicillin) and scored for survival. WT vs *unc-30(ok613),* $P < 0.0001$.

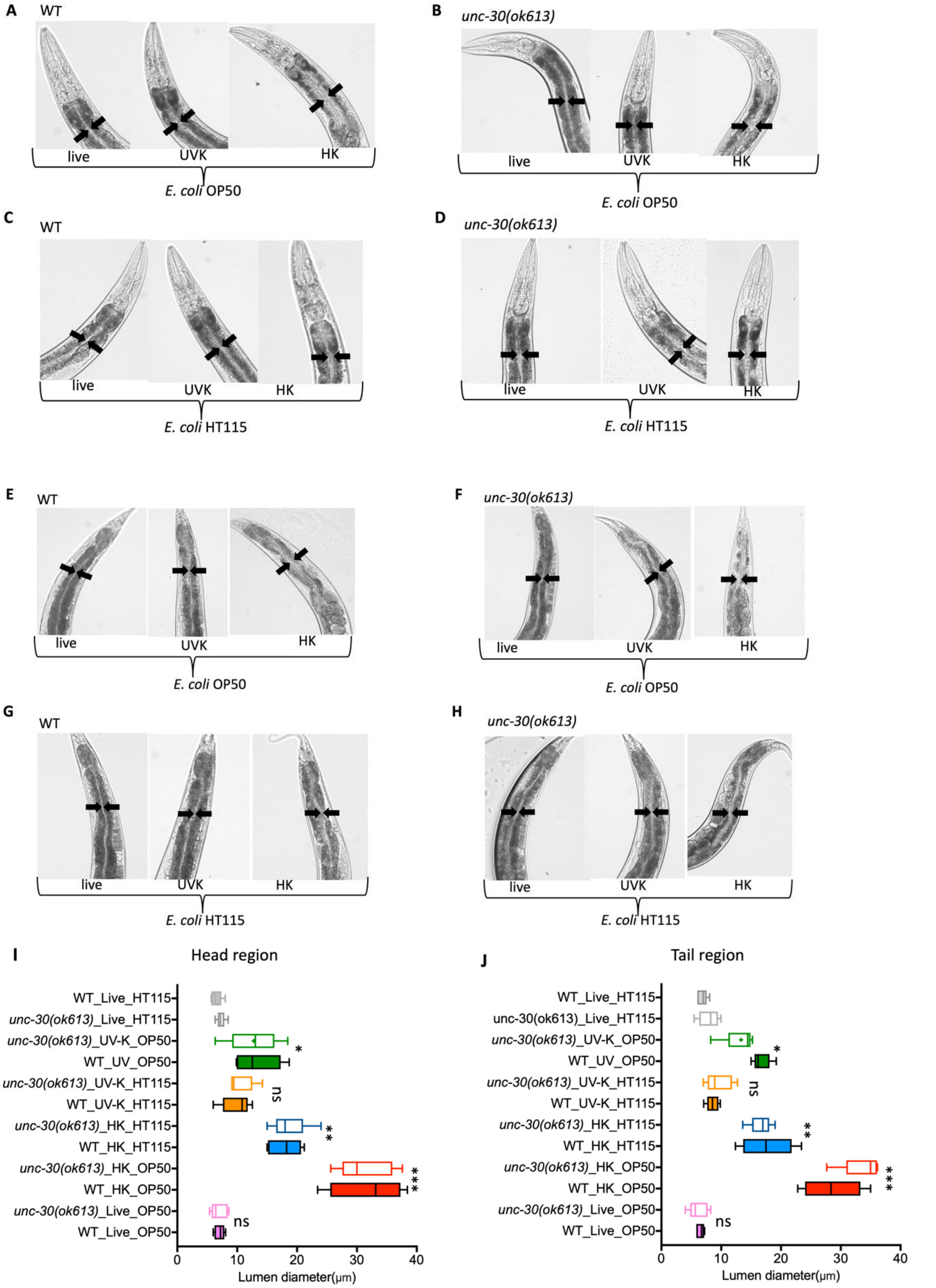


**Figure S2. Effect of heat-killed and UV-killed *E. coli* on *C. elegans* intestinal distension. Related to Figure 1, 4, 5, 6 and S1.**

(A) Representative microscopic images showing anterior intestinal lumen of WT animals on live, UV-, and Heat-killed OP50 (*E. coli*).

(B) Representative microscopic images showing anterior intestinal lumen of *unc-30(ok613)* animals on live, UV-, and Heat-killed OP50 (*E. coli*).
(C) Representative microscopic images showing anterior intestinal lumen of WT animals on live, UV-killed and Heat-, HT115 (*E. coli*).
(D) Representative microscopic images showing anterior intestinal lumen of *unc-30(ok613)* animals on live, UV-, and Heat-killed HT115 (*E. coli*).
(E) Representative microscopic images showing posterior intestinal lumen of WT animals on live, UV-, and Heat-killed OP50 (*E. coli*).
(F) Representative microscopic images showing posterior intestinal lumen of *unc-30(ok613)* animals on live, UV-, and Heat-killed OP50 (*E. coli*).
(G) Representative microscopic images showing posterior intestinal lumen of WT animals on live, UV-, and Heat-killed HT115 (*E. coli*).
(H) Representative microscopic images showing posterior intestinal lumen of *unc-30(ok613)* animals on live, UV-, and Heat-killed HT115 (*E. coli*).
(I) Quantification of the diameter of the anterior intestinal lumen of WT and unc-30(ok613) animals on live, UV-, and heat-killed OP50 and HT115 *E. coli*. Bars represent means while error bars indicate SD; $*p < 0.05$, $**p < 0.001$ and $***p < 0.0001$.
(J) Quantification of the diameter of the posterior intestinal lumen of WT and *unc-30(ok613)* animals on live, UV-, and heat-killed OP50 and HT115 *E. coli*. Bars represent means while error bars indicate SD; $*p < 0.05$, $**p < 0.001$ and $***p < 0.0001$.

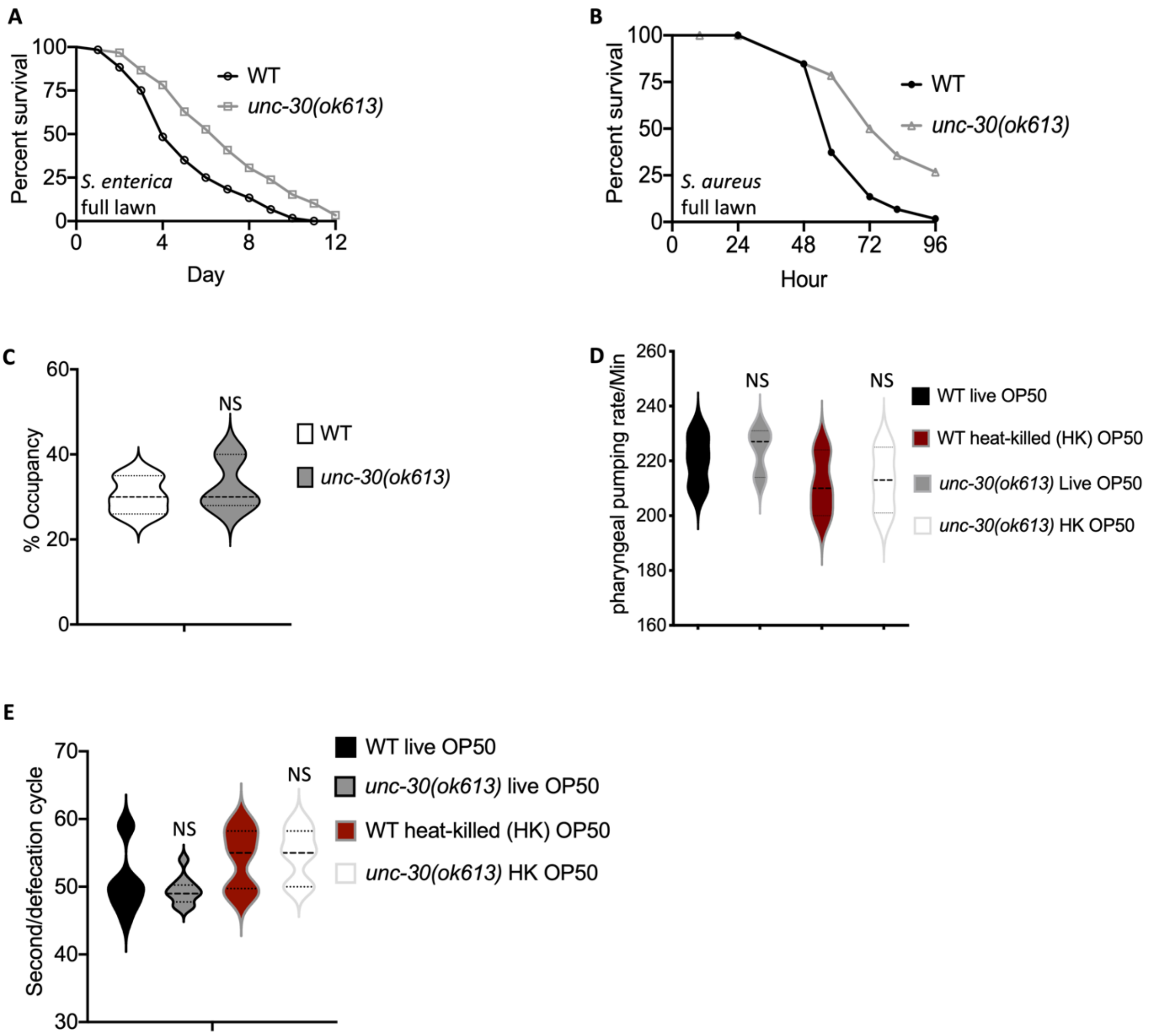


**Figure S3. UNC-30 functional loss enhances *C. elegans* survival against different pathogens. Related to Figure 1.**

(A) WT and *unc-30(ok613)* animals were exposed *S. enterica* full lawn and scored for survival. WT vs *unc-30(ok613)*, P < 0.0001.

(B) WT and *unc-30(ok613)* animals were exposed *S. aureus* full lawn and scored for survival. WT vs *unc-30(ok613)*, P < 0.0001.

(C) WT, *unc-30(ok613)*, and *unc-30(ok613);Prab-3::unc-30* animals lawn occupancy when exposed to partial lawn of culture *P. aeruginosa*. Bars represent means while error bars indicate SD; ***p < 0.0001, P =NS.

(D) Pharyngeal pumping rate of WT and *unc-30(ok613)* animals exposed live and heat-killed *E. coli aeruginosa*. Bars represent means while error bars indicate SD; P =NS.

(E) Defecation cycle of WT and *unc-30(ok613)* animals exposed live and heat- killed *E. coli aeruginosa*. Bars represent means while error bars indicate SD; P =NS.

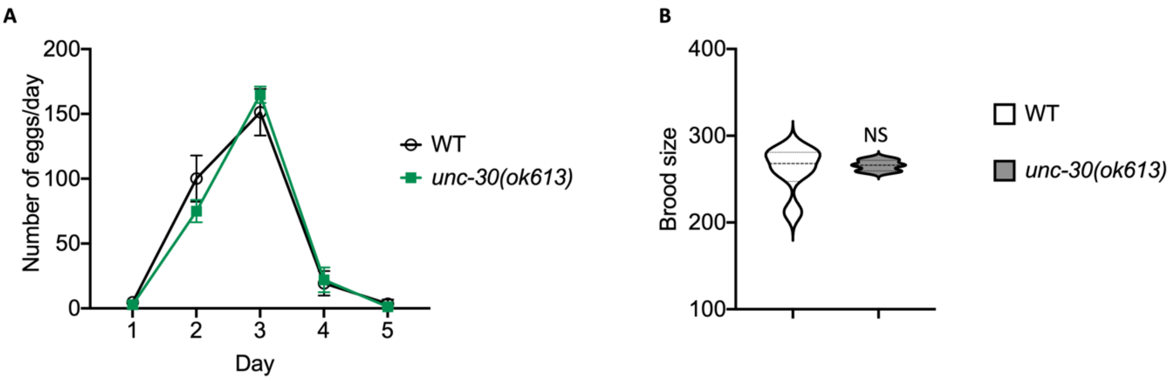


**Figure S4. UNC-30 Functional does not affect the reproductive potential. Related to Figure 1.**

(A) Number of eggs/days for WT and *unc-30*(*ok613*). WT vs *unc-30*(*ok613*), P=NS.
(B) Brood size of WT and *unc-30*(*ok613*). WT vs *unc-30*(*ok613*), P=NS.

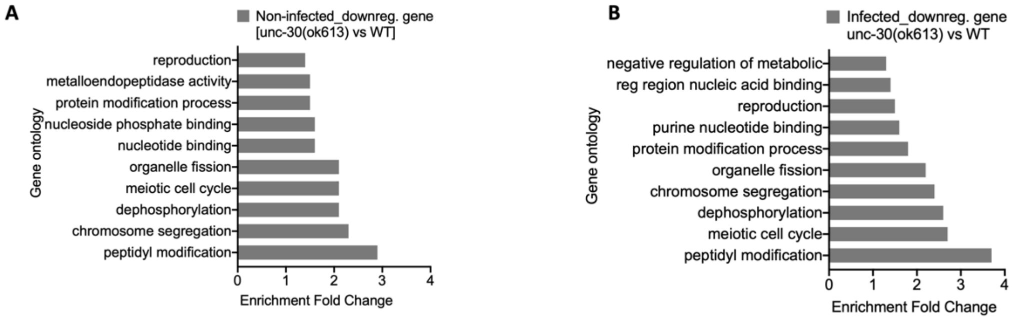


**Figure S5. Transcriptomic enrichment analyses of downregulated genes. Related to Figure 2.**

(A) Enrichment analyses of downregulated genes in non-infected WT and *unc-30*(*ok613*). The cutoff is based on the filtering thresholds of P < 0.05 and arranged according to the representation factor.
(B) Enrichment analyses of downregulated genes in *P. aeruginosa*-infected WT and *unc-30*(*ok613*). The cutoff is based on the filtering thresholds of P < 0.05 and arranged according to the representation factor.

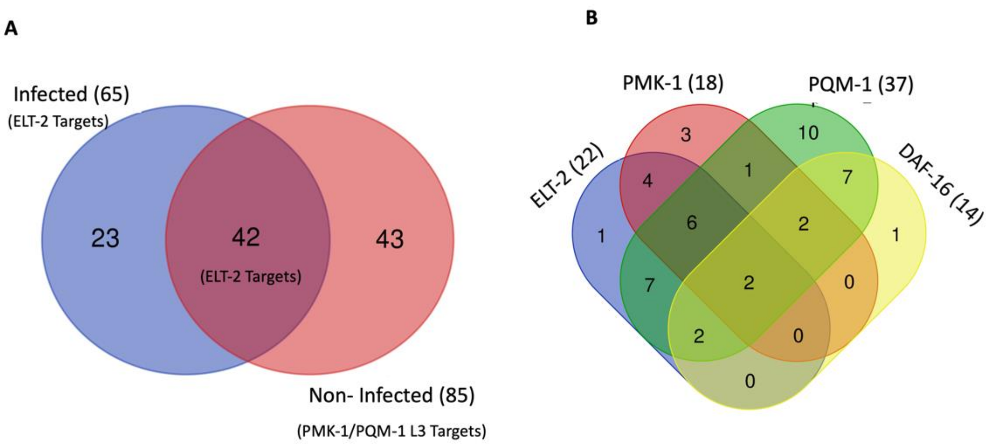


**Figure S6. Enriched pathways regulated by UNC-30. Related to Figure 2.**

(A) Venn diagram of unique and shared immune genes between infected and non-infected animals (*unc-30(ok613)* vs WT).

(B) Venn diagram of unique and shared genes between UNC-30 regulated immune pathways in non-infected animals (*unc-30(ok613)* vs WT).

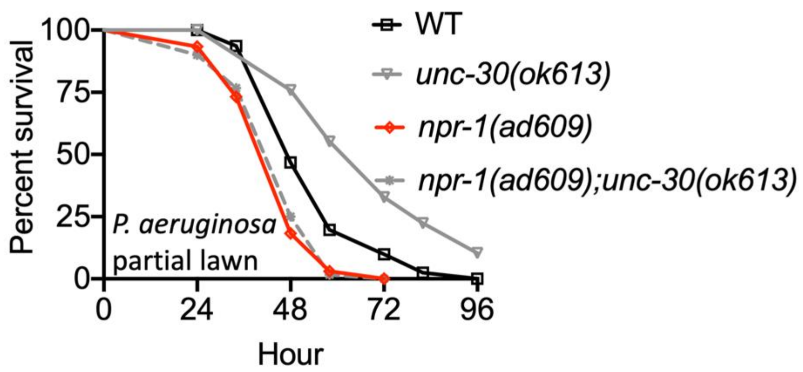


**Figure S7. UNC-30 control immunity via NPR-1. Related to Figure 5.**

WT, *npr-1(ad609)*, *npr-1(ad609)::unc-30(ok613)* and, *unc-30(ok613)* animals were exposed to partial lawn of *P. aeruginosa* and scored for survival. Wild type (WT) vs *unc-30(ok613)*, P < 0.0001; *npr-1(ad609)*, P < 0.001; *npr-1(ad609)::unc-30(ok613),* P <0.05. While *npr-1(ad609)* vs *npr-1(ad609)::unc-30(ok613)*, P = NS.

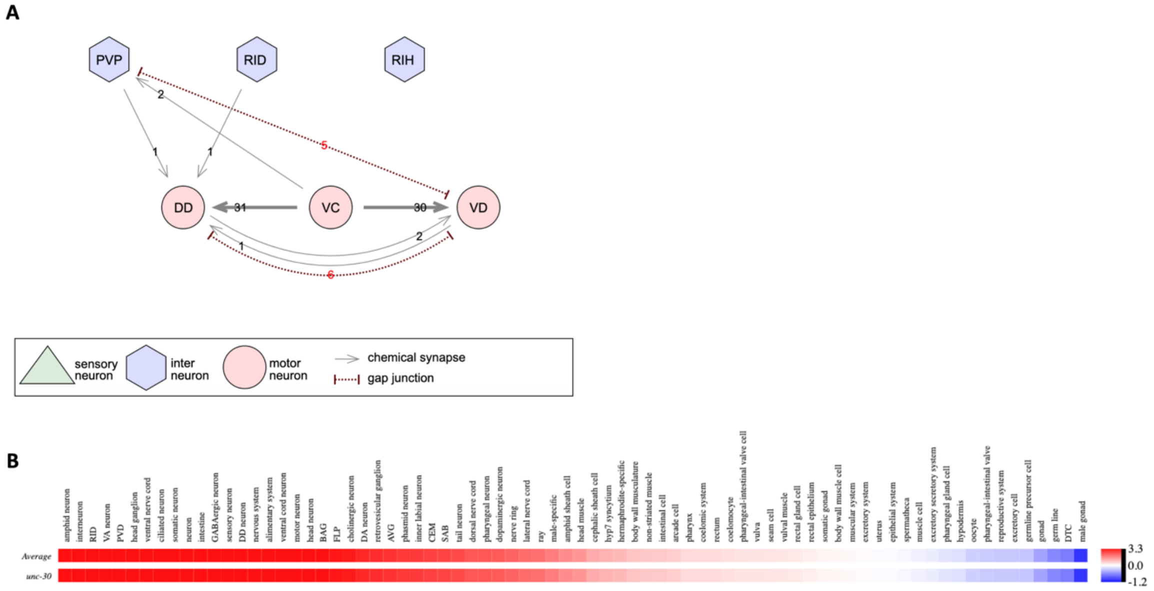


**Figure S8. GABAergic neurons that regulate the expression of UNC-30 in *C. elegans*. Related to Figure 6.**

(A) Neuronal wiring of UNC-30 GABAergic neurons.

(B) Tissue-expression enrichment of UNC-30.

**Table S1. Upregulated and downregulated genes more than 2-folds (p<0.05) in non-infected and *P. aeruginosa*-infected *unc-30(ok613)* vs WT animals. Related to Figure 2.** The RNA seq raw data were submitted and assigned a Gene Expression Omnibus (GEO) accession numbers: GSE149862 and GSE149742.

**Table S2. GO enrichment analysis using Wormbase enrichment tools. Related to Figure 2.**

**Table S3. List of upregulated and downregulated immune genes in non-infected and *P. aeruginosa*-infected *unc-30(ok613)* vs WT animals. Related to Figure 2.**

**Table S4. List of upregulated aging genes and overlapping pathways in non-infected and *P. aeruginosa*-infected *unc-30(ok613)* vs WT animals. Related to Figure 2.**

**Table S5. List of upregulated neuropeptide genes and overlapping pathways in non-infected and *P. aeruginosa*-infected *unc-30(ok613)* vs WT animals. Related to Figure 5.**

**Table S6. Immune pathways from Styer et al., 2008 microarray data analyzed on Worm Exp. Related to Figure 6.**

**Table S7. Mean and maximum survival on different pathogens. Related to Figure 1, 3, 5, 6, S1, and S5.**

**Table S8. Mean and maximum lifespan on live and killed *E. coli*. Related to Figure 1, 4, 5, 6, and S1.**

**Table S9. Primers used in the study. Related to Figures 2, 5 and 6.**
