## Supplementary material for "Immunity-longevity tradeoff neurally controlled by GABAergic transcription factor PITX1/UNC-30": Table S7

Table S7. Mean and maximum survival on different pathogens. Related to Figures 1, 3, 5, 6, S1, and S5.

| **Genotype^1^** | **Bacteria (strain)^2^** | **Mean survival ± SD (Hours)^3^** | **% Change in mean to WT or control)** | **Maximum survival ± SD (Hours) ^3^** | **% Change in maximum to WT or control)** | **N (n) ^4^** | **Figure** |
| --- | --- | --- | --- | --- | --- | --- | --- |
| WT  *unc-30(ok613)* | *P. aeruginosa* (PA14) Full lawn | 52.77**±** 6.94  69.73**±**12.65*** | 56.91 | 83.33**±**12.06  107.33**±**12.06*** | 80.54 | 3(166)  3(149) | Fig. 1A |
| MGH171 EV  MGH171;unc-30 RNAi  *unc-30(ok613)* EV | *P. aeruginosa* (PA14) Full lawn | 50.51**±**4.20  49.43**±**1.49^ns^  72.36**±**3.72*** | -4.14  84.05 | 75.33**±**5.77  86.67**±**8.08*  110.67**±**8.08*** | 42.29  134.86 | 3(138)  3(134)  3(131) | Fig. 1B |
| WT  *unc-30(ok613)*  *unc-30(ok613);Prab-3::unc-30*  *unc-30(ok613);Punc-30::unc-30* | *P. aeruginosa* (PA14) Partial lawn | 51.25**±**1.57  71.10**±**0.67***  51.41**±**1.31 ^ns^  51.59**±**0.70 ^ns^ | 19.85  0.16  0.34 | 106.00**±**0  120.00**±**0***  106.00**±**0 ^ns^  106.00**±**0 ^ns^ | 41.67  0.00  0.00 | 3(177)  3(168)  3(177)  3(173) | Fig. 1G |
| WT  *unc-30(ok613)*  *unc-30(ok613);Prab-3::unc-30*  *unc-30(ok613);Punc-30::unc-30* | *S. enterica*  Partial lawn | 89.28**±**6.21  171.59**±**20.87***  85.56**±**3.12 ^ns^  86.45**±**4.40 ^ns^ | 59.07  0.48  1.00 | 242.25**±**4.5  318.00**±**12***  258.00**±**12 ^ns^  258.00**±**12 ^ns^ | 180.36  33.37  33.65 | 4(227)  4(210)  4(236)  4(234) | Fig. 1H |
| WT  *unc-30(ok613)*  *unc-30(ok613);Prab-3::unc-30*  *unc-30(ok613);Punc-30::unc-30* | *S. aureus*  Partial lawn | 55.28**±**0.19  70.21**±**2.68***  55.62**±**0.84 ^ns^  55.94**±**1.80 ^ns^ | 195.98  8.86  6.73 | 102.67**±**5.77  136.00**±**13.86***  102.67**±**5.77 ^ns^  102.67**±**5.77 ^ns^ | 102.88  0.00  0.00 | 3(176)  3(162)  3(165)  3(180) | Fig. 1I |
| WT  *unc-30(ok613)*  *unc-30(ok613);Prab-3::unc-30*  *unc-30(ok613);Punc-30::unc-30* | *P. aeruginosa* (PA14) Full lawn | 49.85**±**1.89  62.12**±**0.83***  50.73**±**1.48 ^ns^  49.67**±**3.20 ^ns^ | 46.65  1.05  2.05 | 86.67**±**8.08  112.00**±**13.86***  86.67**±**8.08 ^ns^  86.67**±**8.08 ^ns^ | 89.83  0.00  0.00 | 3(158)  3(141)  3(163)  3(128) | Fig. 1G |
| WT  *unc-30(ok613)*  *daf-16(mu86)*  *daf-16(mu86);unc-30(ok613)* | *P. aeruginosa* (PA14) Full lawn | 45.49**±**0.18  62.00**±**0.92***  44.53**±**0.35 ^ns^  58.89**±**1.59*** | 50.96  -2.94  41.38 | 91.33**±**8.08  115.33**±**8.08***  91.33**±**8.08 ^ns^  115.33**±**8.08*** | 74.07  0.00  72.28 | 3(170)  3(162)  3(179)  3(166) | Fig. 3A |
| WT  *unc-30(ok613)*  *pqm-1(ok485)*  *pqm-1(ok485);unc-30(ok613)* | *P. aeruginosa* (PA14) Full lawn | 45.49**±**0.18  62.00**±**0.92***  50.37**±**0.24*  61.58**±**1.85** | 50.96  15.06  49.66 | 45.49**±**0.18  62.00**±**0.92***  50.37**±**0.24*  61.58**±**1.85*** | 59.67  25.00  43.91 | 3(170)  3(162)  3(160)  3(167) | Fig. 3B |
| WT  *unc-30(ok613)*  *elt-2* RNAi  *unc-30(ok613);elt-2* RNAi | *P. aeruginosa* (PA14) Full lawn | 46.47**±**0.70  62.31**±**0.72***  32.80**±**0.98***  32.74**±**0.38*** | 44.73  -38.63  -38.80 | 46.47**±**0.70  62.31**±**0.72***  32.80**±**0.98***  32.74**±**0.38*** | 54.61  -93.33  -91.25 | 3(162)  3(177)  3(175)  3(179) | Fig. 3C |
| WT  *unc-30(ok613)*  *pmk-1(km25)*  *pmk-1* RNAi  *pmk-1(km25);unc-30(ok613)* | *P. aeruginosa* (PA14) Full lawn | 46.47**±**0.70  62.30**±**0.70***  32.22**±**1.12***  31.46**±**0.69***  53.25**±**0.82* | 49.45  -44.55  -46.93  21.17 | 46.47**±**0.70  62.30**±**0.70***  32.22**±**1.12***  31.46**±**0.69***  53.25**±**0.82* | 60.41  -122.48  -112.97  33.53 | 3(170)  3(160)  3(166)  3(180)  3(169) | Fig. 3D |
| WT  *unc-30(ok613)*  *nsy-1(ag3)*  *nsy-1(ag3);unc-30(ok613)* | *P. aeruginosa* (PA14) Full lawn | 45.49**±**0.18  62.00**±**0.92***  35.53**±**0.91***  50.08**±**0.66 ^ns^ | 50.96  -30.74  14.19 | 45.49**±**0.18  62.00**±**0.92***  35.53**±**0.91***  50.08**±**0.66* | 59.67  -93.86  23.95 | 3(170)  3(162)  3(174)  3(167) | Fig. 3E |
| WT  *unc-30(ok613)*  *sek-1(km4)*  *sek-1(km4);unc-30(ok613)* | *P. aeruginosa* (PA14) Full lawn | 46.03**±**0.79  62.08**±**0.85***  31.45**±**0.40***  51.09**±**1.50 ^ns^ | 49.53  -44.99  15.61 | 46.03**±**0.79  62.08**±**0.85***  31.45**±**0.40***  51.09**±**1.50* | 74.07  -118.21  34.14 | 3(170)  3(162)  3(172)  3(166) | Fig. 3F |
| WT  *unc-30(ok613)*  *npr-1(ad609)*  *npr-1(ad609);unc-30(ok613)* | *P. aeruginosa* (PA14) Full lawn | 48.01**±**3.43  67.28**±**4.83***  40.03**±**0.25**  42.94**±**0.17** | 64.65  -25.40  15.27 | 48.01**±**3.43  67.28**±**4.83***  40.03**±**0.25**  42.94**±**0.17** | 69.35  -76.43  -38.15 | 3(145)  3(149)  3(157)  3(166) | Fig. 5C |
| WT  *unc-30(ok613)*  PVP(-) | *P. aeruginosa* (PA14) Full lawn | 54.85**±**5.23  81.11**±**10.83***  56.23**±**3.95 ^ns^ | 83.11  3.36 | 54.85**±**5.23  81.11**±**10.83***  56.23**±**3.95 ^ns^ | 75.94  3.72 | 3(156)  3(158)  3(179) | Fig. 6A |
| WT  *unc-30(ok613)*  ASG(-)  ASG(-)*;unc-30(ok613)* | *P. aeruginosa* (PA14) Full lawn | 45.60**±**1.70  62.39**±**2.71***  59.94**±**2.81**  62.82**±**6.88*** | 49.96  51.20  53.46 | 45.60**±**1.70  62.39**±**2.71***  59.94**±**2.81***  62.82**±**6.88*** | 56.54  67.85  59.00 | 3(164)  3(168)  3(140)  3(161) | Fig. 6B |
| WT  *unc-30(ok613)*  *unc-30(ok613);Pgcy-15::unc-30* | *P. aeruginosa* (PA14) Full lawn | 47.82**±**4.73  61.69**±**1.36***  47.81**±**3.53 ^ns^ | 36.50  -0.03 | 47.82**±**4.73  61.69**±**1.36***  47.81**±**3.53 ^ns^ | 63.15  -10.82 | 3(164)  3(190)  3(154) | Fig. 6C |
| WT  *unc-30(e191)*  *unc-30(e596)*  *unc-30(e318)* | *P. aeruginosa* (PA14) Full lawn | 49.70**±**0.34  63.78**±**2.04***  64.38**±**0.00***  63.67**±**0.00*** | 33.83  32.91  29.59 | 49.70**±**0.34  63.78**±**2.04***  64.38**±**0.00***  63.67**±**0.00*** | 40.86  38.11  36.01 | 4(243)  4(208)  4(223)  4(236) | Fig. S1A |
| WT  *unc-30(ok613)* | *S. enterica*  Full lawn | 87.04**±**5.27  177.59**±**20.92*** | 292.10 | 87.04**±**5.27  177.59**±**20.92*** | 232.25 | 3(169)  3(155) | Fig. S1C |
| WT  *unc-30(ok613)* | *S. aureus*  Full lawn | 55.19**±**0.10  70.29**±**2.61*** | 44.93 | 55.19**±**0.10  70.29**±**2.61*** | 99.20 | 3(178)  3(168) | Fig. S1D |
| WT  *unc-30(ok613)*  *npr-1(ad609)*  *npr-1(ad609);unc-30(ok613)* | *P. aeruginosa* (PA14)  Partial lawn | 48.50**±**2.98  66.05**±**5.99***  40.03**±**0.26**  42.47**±**0.63* | 75.00  -38.85  -25.75 | 48.50**±**2.98  66.05**±**5.99***  40.03**±**0.26*  42.47**±**0.63* | 102.25  -119.27  -111.11 | 3(98)  3(117)  3(109)  3(117) | Fig. S5 |

^1^Genotype: Wild-type hermaphrodites or the indicated mutant/strain were analyzed.

^2^Bacteria: Bacterial species indicated were either spotted at the center on the SK plate to form a partial lawn or spread entirely on the plate to form a full lawn.

^3^Mean, Maximum, % Change: Maximum adult lifespan is the mean lifespan of the 20% of the population for each experiment. Comparisons are to the matched control, which is the WT except for row 2 MGH171 was used as the control. The Kaplan-Meier method was used to calculate the survival fractions, and statistical significance between survival curves was determined using the log-rank test. All error bars represent the standard deviation (SD). The two-sample t-test was used when needed, and the data were judged to be statistically significant when p < 0.05. In the figures, asterisks (*) denote statistical significance as follows: ns, not significant, *, p < 0.05, **, p < 0.001, ***, p < 0.0001, as compared with the appropriate controls.

^4^N: Total number of hermaphrodites analyzed, and the number of independent experiments. Animals were tightly synchronized by egg-laying and allow to develop to young adults at 20ºC and all survival experiments were done with the indicated bacterial at 25ºC. Animals that died due to matricidal hatching were not censored from the data.
