## Supplementary material for "Immunity-longevity tradeoff neurally controlled by GABAergic transcription factor PITX1/UNC-30": Table S8

Table S8. Mean and maximum lifespan. Related to Figures 1, 4, 5, 6, and S1.

| Genotype^1^ | Bacteria^2^ | Mean survival ± SD (Days)^3^ | % Change in mean to WT or control) | Maximum lifespan ± SD (Days) ^3^ | % Change in maximum to WT or control) | N(n) ^4^ | Figure |
| --- | --- | --- | --- | --- | --- | --- | --- |
| WT  *unc-30(ok613)*  *Prab-3::unc-30(ok613)* | Live *E. coli*  HT115 | 13.02±0.64  15.51±0.78***  13.15±0.13 ^ns^ | 13.14  0.63 | 18.00±2.65  20.67±5.86***  20.33±3.21 ^ns^ | 14.11  -1.76 | 3(205)  3(189)  3(195) | Fig. 1D |
| WT  *unc-30(ok613)*  *Prab-3::unc-30(ok613)* | UV-killed  *E. coli*  HT115 | 17.29±0.03  11.84±1.41***  17.35±1.42 ^ns^ | -16.84  0.18 | 28.33±0.58  22.67±0.58***  30.33±0.58 ^ns^ | -17.44  1.03 | 3(324)  3(337)  3(325) | Fig. 1E |
| WT  *unc-30(ok613)*  *mdl-1(tm311)*  *mdl-1(tm311):unc-30(ok613)* | UV-killed  *E. coli*  HT115 | 17.33±0.36  13.38±0.30***  23.22±1.07***  17.64±0.98^ns^ | -12.25  16.69  0.88 | 28.67±0.58  22.33±0.58***  33.33±0.58***  28.67±0.58 ^ns^ | -17.99  13.22  0.00 | 3(323)  3(353)  3(352)  3(331) | Fig.4C |
| WT  *unc-30(ok613)*  *pqm-1(ok485)*  *pqm-1(ok485);unc-30(ok613)* | UV-killed  *E. coli*  HT115 | 17.33± 0.36  13.38±0.30***  18.98±0.25*  17.66±0.44 ^ns^ | -13.23  5.30  0.75 | 28.67±0.58  22.33±0.58***  29.67±0.58 ^ns^  28.67±0.58 ^ns^ | -21.18  24.53  -3.34 | 3(332)  3(350)  3299)  3(310) | Fig.4D |
| WT EV  *unc-30(ok613)* EV  *mdl-1* RNAi  *pqm-1(ok485);unc-30(ok613);mdl-1* RNAi | UV-killed  *E. coli*  HT115 | 17.01±0.61  13.21±0.20***  23.24±1.22***  21.41±0.12** | -8.36  18.76  13.24 | 28.67± 0.58  22.33± 0.58***  33.00±1.00**  33.67± 0.58*** | -13.92  12.94  34.14 | 4(429)  4(477)  4(335)  4(455) | Fig.4E |
| WT EV  *unc-30(ok613)* EV  *skn-1 RNAi*  *unc-30(ok613);skn-1* RNAi | UV-killed  *E. coli*  HT115 | 17.01±0.61  13.21±0.20***  16.39±2.10*  11.69±0.13*** | 8.91  -3.96  -1.41 | 29.33±0.58  22.33±0.58***  27.33±0.58**  23.33±0.58*** | -16.39  -4.68  -14.05 | 4(427)  4(470)  4(442)  4(210) | Fig.4F |
| WT  *unc-30(ok613)*  *npr-1(ad609)*  *npr-1(ad609);unc-30(ok613)* | UV-killed  *E. coli*  HT115 | 17.33±0.36  13.38±0.30***  16.93±0.10 ^ns^  17.38±0.89 ^ns^ | -11.30  -1.19  0.14 | 28.67±0.58  22.33±0.58***  29.67±0.58 ^ns^  29.67±0.58 ^ns^ | -18.52  21.44  0.00 | 3(332)  3(350)  3(342)  3(326) | Fig.5D |
| WT  *unc-30(ok613)*  ASG(-)  PVP(-) | UV-killed  *E. coli*  HT115 | 17.33±0.36  13.38±0.30***  22.53±0.51***  17.00±0.23 ^ns^ | -11.60  15.24  -0.99 | 28.67±0.58  22.33±0.58***  33.67±0.58***  28.67±0.58 ^ns^ | -18.57  33.24  0.00 | 3(332)  3(350)  3(341)  3(341) | Fig. 6E |
| WT  *unc-30(ok613)*  ASG(-)  ASG(-);*unc-30(ok613)* | UV-killed  *E. coli*  HT115 | 17.33±0.36  13.38±0.30***  22.52±0.51***  21.25±0.61*** | -11.60  15.82  12.10 | 28.67±0.58  22.33±0.58***  33.67±0.58***  33.33±0.58*** | -18.57  33.24  0.00 | 3(332)  3(350)  3341)  3328) | Fig. 6F |
| WT  *unc-30(ok613)*  ASG(-)  *unc-30(ok613);Pgcy-15::unc-30(ok613)* | UV-killed  *E. coli*  HT115 | 17.33±0.36  13.38±0.30***  22.52±0.51***  16.62±1.36 ^ns^ | -11.92  16.17  -2.35 | 28.67±0.58  22.33±0.58***  33.67±0.58***  31.00±1.00* | -19.09  34.14  1.00 | 3(324)  3(337)  3(332)  3(321) | Fig. 6G |
| WT  *unc-30(ok613)* | live  *E. coli*  OP50::GFP | 8.99±0.55  13.93±1.06*** | 33.84 | 20.50±0.71  24.50±0.71*** | 27.40 | 3(173)  3(147) | Fig. S1B |
| WT  *unc-30(ok613)* | UV-killed  *E. coli*  HT115 | 13.43±0.52  10.79±0.40*** |  | 19.50±0.71  17.00±1.41*** |  | 3  3 | Fig. S1D |

^1^Genotype: Wild-type hermaphrodites or the indicated mutant/strain were analyzed.

^2^Bacteria: Bacterial species of the indicated strains were spotted in the center of the NGM dish to form a small lawn. To generate dead *E. coli*, we spotted bacteria on the dish and then treated with UV light.

^3^Mean, Maximum, % Change: Maximum adult lifespan is the mean lifespan of the 20% of the population that had the longest lifespans. Comparisons are to the matched control, which is the WT, control. The Kaplan-Meier method was used to calculate the survival fractions, and statistical significance between survival curves was determined using the log-rank test. All error bars represent the standard deviation (SD). The two-sample t-test was used when needed, and the data were judged to be statistically significant when p < 0.05. In the figures, asterisks (*) denote statistical significance as follows: ns, not significant, *, p < 0.05, **, p < 0.001, ***, p < 0.0001, as compared with the appropriate controls.

^4^N: Total number of hermaphrodites analyzed, and the number of independent experiments. Animals were tightly synchronized by egg-laying and allow to develop to young adults at 20ºC and all survival experiments were done with the indicated bacterial at 20ºC, expect it is indicated.
