## Supplementary material for "Immunity-longevity tradeoff neurally controlled by GABAergic transcription factor PITX1/UNC-30": Table S9

| **Primer for quantitative reverse transcription-PCR** | | |
| --- | --- | --- |
| **Gene name** | **Forward primer sequence (5'-3')** | **Reverse primer sequence (5’-3’)** |
| *Pan-act* | TCGGTATGGGACAGAAGGAC | CATCCCAGTTGGTGACGATA |
| *Unc-25* | CCCTCTTTGAGCAAGGGTTTA | CTGTGGCTCAGTTGTGTAGTT |
| *Unc-47* | GGAGAAGCATCAGAGCCAATA | CCACCATCCACCAACCTTTA |
| *Lys-7* | CCTTTGCCTGTCTCCCTTAAT | CCTTTGCCTGTCTCCCTTAAT |
| *K12H4.7* | GGAGCCTGGGTCTTTGATATT | TAGCTTGGGCAGAAGACAAG |
| *lys-8* | CTCCACGAGTTCCACCAAAT | CGGACAAAGACTGCCGAATA |
| *Irg-5* | CGGTGAAGAAGTACGACCAATA | GTGTACCATGCATCAGCATTATC |
| *Lys-4* | CTTCCCATGCTTGAGATCTAACT | AGCCAGAGAGTAGAGACATGAG |
| *Lys-1* | TCGGAATCTACACCAACCAATAC | GTAACACCACCTCCAAGAACA |
| *nlp-21* | TGTGGAGAACTCGAAGAGAGA | TGGCTTGACCGGAAATGAA |
| *nlp-35* | GCGCTTCAGAAGATCCGTATAA | GGGAAGTAAGCCACTAACAACA |
| *nlp-22* | CTCCGAACCCTTCCAATCAT | CCGATTGGGAATCCAGTTAGT |
| *flp-21* | CGTCCTCTCCGATTTGGATAAA | GTGGAGTGTCTGGAACATCAT |
| *flp-18* | TCGGGCGTTCTTCTCATTTC | CCTCTGCTGGTATGTCTTCTTC |
| *flp-7* | AGAACCAACTGAACTCGAGAAG | GTCCGAACCGAACCATTGA |
| *flp-5* | CTTCATCGCCACCCTTCTT | TTTGGTGCTCGTGCTATTCT |
| *rps-6* | GTCAGAATCGGAGGAGGAAAC | GCAGGATTGTCCCTTCTTCA |
| *rpl-19* | GGTACTGCC AATGCTCGTAT | GAAGGTAGAGCTCGTGGTAAAG |
| *rpl-30* | TGGTCATGAAGACTGGACAATAC | TGAGTGGTGGAGTGTTGTTG |
| *atp-3* | TCCCAAGCAAGCCTCTTTAC | CTCGACTCCATGAACCTGAATC |
| *rpl-9* | GATCCGTAACTTCTTGGGAGAG | GGACATCATTTCCCTCGACTAC |
| *mdl-1* | TCAAGCCGACGAAGAAGTTG | CGAGAAGATGTGGAGGTGAAAG |
| **Primer for generation of transgenic animal** | | |
|  | **Forward primer sequence (5'-3')** | **Reverse primer sequence (5’-3’)** |
| *unc-30* | ATGCGAAATCAGTTGAGAATTCTGCG | GTGTTGATCTTTTGCCGGAGGG |
| *Ptax-2* | ATCAGCATCAGCTGTCGTATCTCC | ATCGGAAAACTCCGGTTTTTCTGACA |
| *Plim-6* | GAAATTATCAAGCATACTGCGGTACATCAAAAAA | CCCTGTAGATATACCGATCTTTGATAAGACACC |
| *Podr-2* | CCTATTAGAACATTGAACCAATTCT | CGGGCATCCCGACAAACT |
| *Punc-53* | CACTTGGTGTCAGAGTATCCCATTT | TTTATTCGGAGCACCAACATTTCGT |
| *Pgcy-15* | TATATTGTGGCCATAGCGTCATTTCCT | AGCTGATGGGATGTAGGCAGC |
| *Pgcy-15_unc-30* | TACATCCCATCAGCTATGGATGACAATACGGCCACAC | TATGGCCACAATATACTAAAGTGGTCCACTGTACTGAC |
